# LHX2 in germ cells control tubular organization in the developing mouse testis

**DOI:** 10.1101/2022.12.29.522214

**Authors:** Neha Singh, Domdatt Singh, Anshul Bhide, Richa Sharma, Shilpa Bhowmick, Vainav Patel, Deepak Modi

**Affiliations:** Molecular and Cellular Biology Laboratory, ICMR-National Institute for Research in Reproductive and Child Health, Indian Council of Medical Research (ICMR), JM Street, Parel, Mumbai 400012, India; Viral Immunopathogenesis Laboratory, ICMR-National Institute for Research in Reproductive and Child Health, Indian Council of Medical Research (ICMR), JM Street, Parel, Mumbai 400012, India

**Keywords:** Homeobox, LIM-HD gene, Knockout, Sex determination, Gonad, development, Interstitial cell, Endothelial cell, Infertility, Male development, Seminiferous tubules

## Abstract

In the gonads of mammalian XY embryos, the organization of cords is the hallmark of testis development. This organization is thought to be controlled by interactions of the Sertoli cells, endothelial and interstitial cells with little or no role of germ cells. Challenging this notion, herein we show that the germ cells play an active role in the organization of the testicular tubules. We observed that the LIM-homeobox gene, *Lhx2* is expressed in the germ cells of the developing testis between E12.5-E15.5. In *Lhx2* knockout-fetal testis there was altered expression of several genes not just in germ cells but also in the supporting (Sertoli) cells, endothelial cells, and interstitial cells. Further, loss of *Lhx2* led to disrupted endothelial cell migration and expansion of interstitial cells in the XY gonads. The cords in the developing testis of *Lhx2* knockout embryos are disorganized with a disrupted basement membrane. Together, our results show an important role of *Lhx2* in testicular development and imply the involvement of germ cells in the tubular organization of the differentiating testis.

**Highlight:**

- *Lhx2* is expressed in germ cells of developing mouse testis but is dispensable for germ cell and Sertoli cell specification
- Loss of *Lhx2* disrupts testicular vascularization, leads to the expansion of interstitial cells, and alters the tubular organization
- Germ cells govern the tubular organization in the developing testis

**Graphical Abstract:** 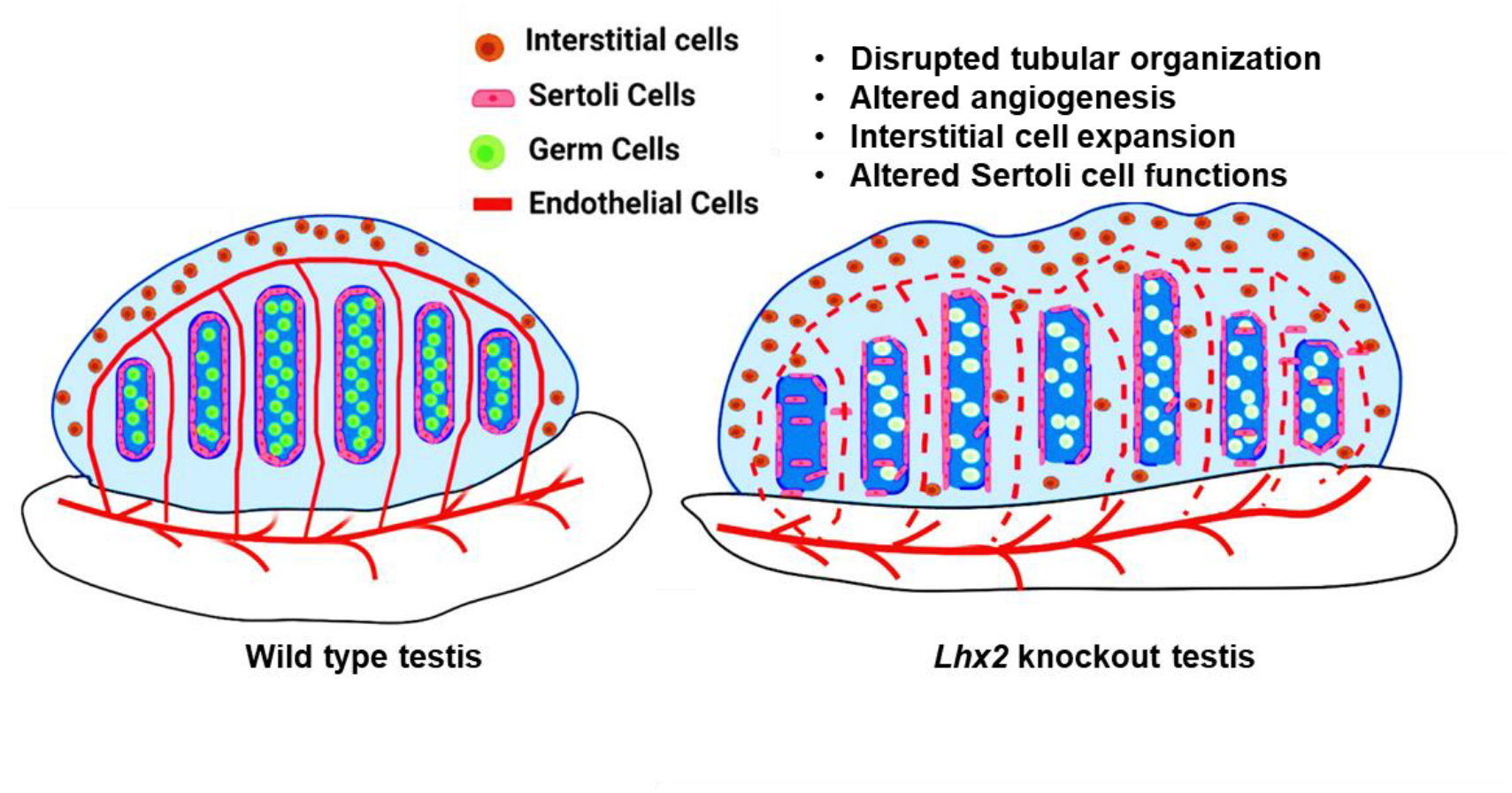

## Introduction

Mammalian sex determination is characterized by the differentiation of the bipotential gonad into the testis or an ovary [1]. In the XY embryos, testicular development requires Sertoli cell specification and tubular organization, vascularization, and differentiation of the interstitial cells to Leydig cells, peritubular myoid cells, and other cell types. In the XY embryos, SRY-positive precursors differentiate into Sertoli cells that act as a scaffold for the germ cells to aggregate, organize and start forming the testicular cords [2,3]. The number of Sertoli cells is the key determinant of tubulogenesis and reduced numbers of the Sertoli cells compromise tubular organization [4].

The tubular organization is further supported by the migrating vascular endothelial cells from the adjacent mesonephros [5]. Blocking mesonephric cell migration in developing testis either directly by a barrier [6,7] or by inhibiting VE-Cadherin [8,9] leads to defective cord formation. Sertoli cells secrete a plethora of factors that activate multiple signal transduction events in the endothelial cells to aid in their migration from the mesonephros [10–12]. Inhibition of the PI3K signaling pathway [13], PDGF signaling [14,15], or VEGF signaling [8,16–18] disrupts the organization of tubules in the developing testis. These observations imply the involvement of Sertoli and endothelial cell communication for the tubular organization.

In addition to endothelial cells, interstitial progenitors, fetal Leydig cells, peritubular myoid cells, and macrophages also aid in tubule organization. Blocking the macrophage migration into the testis [9], genetic disruption of gene encoding activin A from fetal Leydig cells [19], and defective or reduced numbers of peritubular myoid cells [20–22] result in altered tubular morphology. These results imply the involvement of interstitial cells in the structural organization of tubules in testis development.

Somatic cells have an unequivocal role, while the role of germ cells (if any) in the tubulogenesis organization is underexplored. A single study has demonstrated that fetal mice lacking germ cells (due to a point mutation in the cKIT gene) have defective tubule organization due to compromised Sertoli cell functions [23]. However, it is unclear if the tubules were hypoplastic due to the physical absence of the germ cell core or due to the absence of some factors secreted by the germ cells. Thus, the involvement of germ cells in the tubular organization is unclear.

Lim homeobox gene 2 (*Lhx2*), is a transcription factor of the LIM-HD family, having roles in brain patterning, eye development, olfaction path, hair folliculogenesis, limb bud outgrowth, and several others [24,25]. Though the functions of *Lhx2* are well-reported in the developing nervous system, its expression and function in the developing gonads is not explored. Members of the LIM-HD family are expressed in the developing gonads of multiple species including humans, of them the roles of *Lhx1, Lhx8*, and *Lhx9* are well reported [26,27]. Recently we reported that the Lim-HD gene *Lhx2* is expressed in the germ cells of the developing XX gonads, and loss of *Lhx2* promotes endothelial cell migration in the developing ovary [26]. To our knowledge, this is the first study demonstrating the involvement of germ cells in controlling endothelial cell migration and vascularization. Whether *Lhx2* is also expressed in the germ cells of XY gonads and if it governs vascularization is unknown. Further, the functions of *Lhx2* in the developing gonads beyond the suppression of vascularization are not known.

In the present study, we aimed to investigate the expression of *Lhx2* and its function in developing testis. The results revealed that *Lhx2* is expressed by germ cells of the fetal testis and loss of *Lhx2* leads to disrupted tubular organization. A preprint version of this manuscript is available [28].

## Materials and Methods

### Animal Ethics, Mice Strain and nomenclature

The study was approved by the institutional animal ethics committee (IAEC Project No., 27/14, 2/16, 16/16). *Oct4*-GFP transgenic mice [29] and cKit^W^/^+^ mice [30] were kindly provided by Dr. K.V.R. Reddy, NIRRCH. FVB mice constitutively expressing Green fluorescent protein (GFP) were provided by the National Institute of Immunology (NII), New Delhi, India. Heterozygous mice with a targeted disruption of *Lhx2* [31] and those with conditional *Lhx2* allele [32] were kindly provided by Prof. Shubha Tole, TIFR, Mumbai, India. All the animals were bred and maintained at the Experimental Animal Facility of the Institute.

In the present study, pups of pregnant dams with *Lhx2* flox alleles along with tamoxifen-inducible CreER and treated with tamoxifen are referred to as *Lhx2*^*flox/flox*^. Pregnant dams with *Lhx2* flox alleles without the tamoxifen-inducible CreER and treated with tamoxifen are labeled as wild-type (WT) controls. In the case of standard *Lhx2* knockouts, the homozygous knockout pups are referred to as *Lhx2*-/-(or mutants) and the homozygous wild-type pups as WT controls. For the germ cell-depleted mice, the cKit^+^/^+^ are WT and cKit^W^/^W^ are homozygous mutants.

Animals were mated and the day of vaginal plug was considered as 0.5; the pups were collected from E11.5 to E15.5 as per experimental requirement. The gonads were dissected under a stereomicroscope (Olympus; Tokyo, Japan); the mesonephros was removed and the gonad was stored in RNALater for RNA extraction. For histology and immunolabeling experiments, the whole pups or isolated gonads were fixed overnight in 4% paraformaldehyde and processed for paraffin embedding and sectioning.

### Genotyping

Primer sequences for all the genotyping are mentioned in Supplementary Table 1. Genotyping was done using the Terra™ PCR Direct kit (Takara Bio, Kusatsu, Shiga Japan). Embryonic sex and *Lhx2* allele genotyping were performed as mentioned [26]. For germ cell-depleted gonads, the embryos from cKit^W^/^+^ crosses were collected and genotyping was done by PCR [30]. In brief, the PCR product of 231 bp was digested with Hph1 restriction enzyme and resolved on 15% polyacrylamide gel. cKIT^W/W^ mutant pups lacked fragments of 40 bp and 70 bp bands while homozygous WT lack a fragment of 111 bp.

### RNA extraction, cDNA synthesis, and real-time PCR

RNA extraction was done using RNeasy mini kit (Qiagen, Germany). cDNA was synthesized using High Capacity cDNA synthesis kit (Applied Biosystems, ThermoFisher Scientific, MA, USA) as mentioned earlier [26]. Real-time PCR was performed using Eva Green chemistry (BioRad Laboratories Inc., CA, USA) in the CFX96 Real-Time PCR System (BioRad) as detailed previously [26]. PCR was carried out in duplicates, for all biological replicates. Gene expression was normalized to *Gapdh* and data were analyzed using the Pfaffl method [33]. The primer sequences, their annealing temperatures, and expected product sizes are given in Supplementary Table 1.

### Section *In Situ* hybridization

Section *in situ* hybridization was performed on 7 µm thick paraffin sections of wild-type fetal testes at E13.5 as detailed previously [34,35]. In brief, deparaffinized sections were acetylated, treated in proteinase-K, and pre-hybridized at 42^0^C. The plasmids for *Lhx2* sense and antisense probe preparation were a kind gift from Prof. Shubha Tole [36]. Digoxigenin (DIG)-labeled sense and antisense riboprobes were prepared using the *in vitro* transcription DIG labeling kit (Roche, IN, USA). Hybridization was carried out at 60^0^C overnight and the slides were washed and probed with an alkaline phosphatase-conjugated anti-DIG antibody (Roche) at 1:2000 dilution. The slides were washed and detection was performed at pH 9.5 by incubating in nitro blue tetrazolium and 5-bromo-4-cholro-2-indoyl phosphate solution (Roche) as per manufacturer instructions. The reaction was stopped, the slides were dehydrated, air-dried, and mounted. The slides were observed under a brightfield microscope (Olympus) and representative areas were photographed.

### Fluorescence-Activated Cell Sorting (FACS) from *Oct4*-GFP testis

XY gonads from *Oct4*-GFP transgenic pups were collected on E13.5 and E15.5 and cell sorting was done [37]. Briefly, the gonads were incubated in 0.25% trypsin-EDTA (Gibco, Thermo Fisher Scientific, MA, USA) at 37^0^C for 20 min and the reaction was stopped by adding an equal volume of DMEM medium supplemented with 10% fetal bovine serum (both from Gibco), followed by mechanical disruption. The cell suspension was filtered with 40µm strainer (BD Bioscience, San Jose, CA) and stained with 1µg/ml concentration of propidium iodide (Sigma-Aldrich) for live/dead discrimination. Fluorescence-activated cell sorting was performed using 70µm nozzle based on GFP signal on a BD FACS Aria fusion Flow Cytometer (BD Biosciences). The GFP positive (germ cells) and negative (somatic cells) live cells (propidium iodide negative) were sorted (Supplementary Figure 1A), and subjected to RNA extraction, cDNA synthesis, and qPCR as described above.

### Histology and Immunohistochemistry/Immunofluorescence

Hematoxylin-Eosin staining and immunohistochemistry were performed on 5 µm thick paraffin sections as mentioned [38,39]. For immunohistochemistry, antigens were retrieved by boiling the sections in Tris-EDTA Buffer (pH 9), followed by blocking in 5% bovine serum albumin (Sigma-Aldrich). Sections were probed overnight with primary antibodies. Negative controls were incubated either with isotype controls or Phosphate Buffered Saline instead of the primary antibody. Detection was done using the Horse Radish Peroxidase (HRP) streptavidin method using ABC staining system (Santa Cruz Biotechnology; Texas, USA). All sections were briefly counterstained with Mayer’s hematoxylin and mounted in DPX liquid mountant (Himedia, Maharashtra, India). Slides were viewed under a brightfield microscope (Olympus) and representative areas were photographed using a digital camera (Olympus). For VE-cadherin, the signals were detected using a donkey anti-rabbit secondary antibody conjugated with Alexa Fluor 568 dye (Invitrogen, A10042; 1:1000 dilution) as described previously [40].

For multiplex immunofluorescence, we used the Tyramide signal amplification (TSA) detection system (PerkinElmer, MA, USA, or Biotium, Inc. CA, USA) as per the manufacturer’s protocol. In brief, the sections were deparaffinized, rehydrated, and re-fixed in 10% neutral buffered formalin. Antigen retrieval was done in Tris-EDTA buffer (pH 9). Slides were washed in 1X tris-buffered saline with 0.1% Tween-20 followed by incubation in blocking solution for 10 min at room temperature. Sections were probed with primary antibody against (anti-DDX4 for germ cells), for 30 min at room temperature. Incubation with anti-rabbit-secondary antibody tagged with HRP was done for 10 min and signal amplification was performed using fluorophore 570. The antibody complex was stripped in boiling in antigen retrieval treatment for 30 minutes, blocking was done and sections were probed with anti-SOX9 antibody (for Sertoli cells) for 30 min at room temperature. Downstream steps were repeated and fluorophore 690 was used to amplify the signals. Eventually, sections were mounted with EverbriteTM hardest mounting medium (Biotium). Laminin staining was done using the same procedure except that the detection was done using the fluorophore 520. Details of all the antibodies and their optimized concentrations used are given in Supplementary Table 2.

Slides were viewed and images were captured using a fluorescence microscope equipped with an sCMOS camera (Leica Microsystems DMi8, Mannheim, Germany). The contrast was enhanced digitally but raw images were used for the quantification (Fiji version of ImageJ).

### RNA Sequencing

The strategy of gonad collection from *Lhx2*^*flox/flox*^ pups for RNAseq is detailed previously [26]. A single dose of tamoxifen (75µg/g) was given orally to gravid females on E9.5 and gonads were collected on E12.5 from the WT and *Lhx2*^*flox/flox*^ embryos. This concentration and time were chosen as they gave maximum efficiency of knockout [26,32]. Approximately, 30 pairs of gonads were pooled, RNA extracted, and outsourced for RNA sequencing on the Illumina NextSeq 500 platform (Sandor Biosciences, Hyderabad, India). An average of, 45 million raw reads were generated per sample. Alignment was done using Tophat 2.0.13 software and mm84 sequence as reference (ftp://ftp.ensembl.org/pub/release-84/fasta/mus_musculus/dna/).

### Analysis of differentially expressed genes in cell-type-enriched transcriptome of the developing testis

Genes with fragments per kilobase per million (FPKM) values >0.5 were selected and the fold change between the *Lhx2*^*flox/flox*^ and WT was calculated. Genes with fold change ≤0.5 were taken as downregulated whereas those with fold change ≥1.5 were taken as upregulated. The list of genes enriched in the supporting cells, germ cells, endothelial cells, and interstitial cells were retrieved from the dataset GSE27715 [41] and the FPKM values from the WT and *Lhx2*^*flox/flox*^ testis were compared. The fold change values were calculated as above. The differentially expressed genes for the total and cell type-specific transcriptomes were subjected to gene ontology analysis using DAVID (https://david.ncifcrf.gov/tools.jsp). In addition, the expression profile of LIM-HD co-regulators in the different cell types of developing testis during the window of sex determination was determined from the dataset GSE27715 [41].

### Gonad Recombination Assay and assessment of cell migration

The scheme of the gonad recombination assay is described in detail previously [26]. In brief, the gonads and mesonephros were carefully dissected from WT and *Lhx2-/-*. XY embryos at E12.5. The dissected gonad was carefully overlayed onto the donor mesonephros from GFP XY embryos of the same gestational age and the assembly was cultured at 37^0^C. The. Recombinants were imaged for the migration of the GFP-positive cells at 24, 48, and 72 hours post-recombination.

Quantification of angiogenic patterns was done using the angiogenesis analyzer macro in ImageJ software as detailed previously [26]. In brief, the background of desired areas was corrected and the images were subjected to the angiogenesis analyzer (ImageJ) that quantified various branching parameters.

### Imaging, Cell number counting, and Morphometric Analysis

Quantification of all signal intensities in immunofluorescence images was done using the LasX software (Leica Microsystems) or ImageJ (https://ij.imjoy.io/). For quantification, 5 random areas were selected per section and 3 non-serial sections per animal were analyzed.

For quantifying tubule numbers and abnormal tubules, five hematoxylin and eosin-stained testis sections (each 20µm apart) were analyzed. The total number of tubules was counted in each section and the number was divided by the area of the section. Tubules appearing close to each other without a clear interstitium were defined as fused tubules. Incomplete tubules were those which did not have well-defined boundaries and appeared interspersed with the stroma. Histologically well-defined tubules without an outer layer of cells were defined as tubules without peritubular myoid cells.

The numbers of DDX4-positive and SOX9-positive cells per testis cord were counted using Image J software. For overall cell number, the putative tubule area was marked using the region of interest (ROI) tool and the number of SOX9 or DDX4 positive cells in that area were quantified in single-channel images. Three to five non-serial sections per animal were quantified.

ImageJ was used to measure the length (the longest diameter), width (the shortest diameter), and rete testis length (testicular region attached to mesonephros) from WT and mutant testis.

### Statistical analysis and data presentation

Data were analyzed statistically using an unpaired Student-t-test or multiple t-test with Holm-Sidak corrections considering random distribution and unequal variance. All results were presented as mean ± standard deviation (SD) and graphs were plotted on GraphPad Prism (Version 8, CA) and Microsoft Excel.

## Results

### *Lhx2* is expressed in germ cells of developing mouse testis

Fig. 1A shows the quantification of *Lhx2* mRNA in the testis of wild-type (WT) embryos from E11.5 to E15.5 (n=6 testis/time point). *Lhx2* transcripts were detected in all the samples irrespective of the gestational age, their levels were highest at E11.5 which gradually and significantly declined at all the time points tested. To determine the cellular localization of *Lhx2*, RNA *in situ* hybridization (n=5) and immunohistochemistry (n=7) were performed on fetal testis sections at E13.5. *Lhx2* transcripts (Fig. 1B) were localized in cells within the cords of developing testis; the interstitium and the celomic epithelium were devoid of any signals indicating that *Lhx2* is either in the developing Sertoli cells or in the germ cells. No signals were detected in the sections incubated with the sense probe. Immunohistochemical analysis (Fig. 1C) revealed that LHX2 was localized in large-size germ cells (open arrow) enclosed within the testis cords. SOX9, a positive control, stained the Sertoli cells around the germ cells (open arrow). No staining was detected in the negative control sections. These results implied that *Lhx2*/LHX2 may be expressed in the germ cells.

**Fig. 1:**
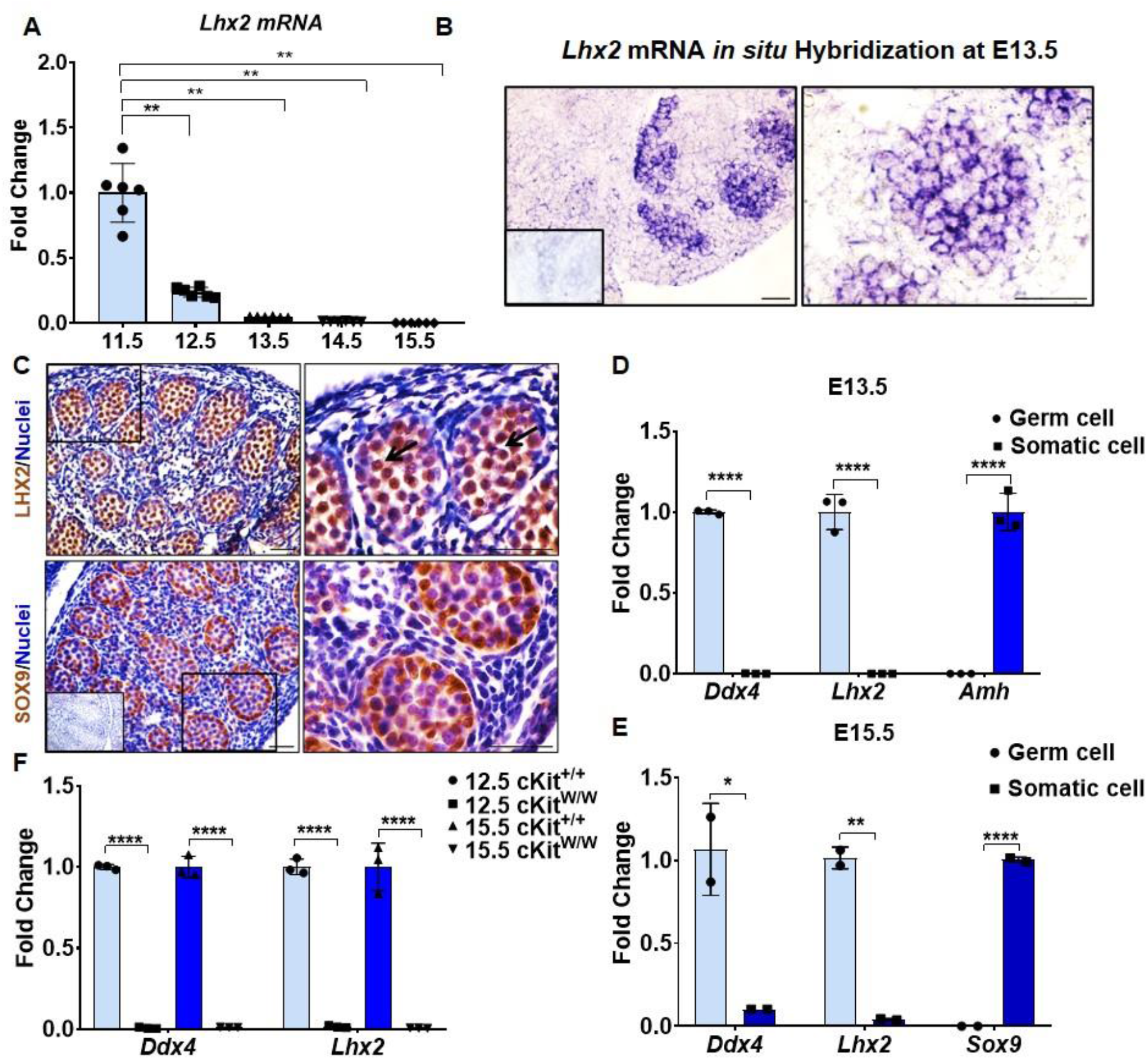
*Lhx2* is expressed in germ cells of developing mouse testis. (A) mRNA expression of *Lhx2* in XY gonads at E11.5-E15.5. Data is presented as mean ± SD (n=6 testis/timepoint). Y axis is fold change of normalized expression value where the mean of E11.5 was taken as 1. (B) Section *in situ* hybridization for *Lhx2* at E13.5 testis (n=5). Purple staining is a positive reaction. Inset is sense probe control. Bar is 50µm. (C) Immunostaining for LHX2 in XY gonads at E13.5 (n=7). SOX9 is a positive control. Arrowheads indicate germ cells. Inset is negative control without primary antibody. Bar is 50µm. Detection of *Lhx2* mRNA in flow-sorted germ cells and somatic cells of XY gonads at E13.5 (D) & E15.5 (E). Germ cells were flow-sorted from XY mice expressing GFP under the *Pou5f1* promoter. Germ cell and somatic cell fractions from several mice at E13.5 and E15.5 were pooled and qPCR for *Ddx4* (germ cell marker), *Amh/Sox9* (somatic cell markers), and *Lhx2* was carried out for three independent batches at E13.5 and two batches at E15.5. Y axis is fold change of normalized relative expression where the mean of the data for germ cell fraction was taken as 1 (for *Ddx4* and *Lhx2*) and somatic cell fraction was taken as 1 for *Amh/Sox9*. (F) qPCR for *Ddx4* and *Lhx2* in XY gonads of cKit knockout (cKit^W/W^) mice at E12.5 and E15.5 (n=3 /timepoint/genotype). Data is presented as mean ± SD. Y axis is the fold change of normalized expression values where the mean value of WT is taken as 1. In all qPCR experiments, *Gapdh* was used for normalization. *, **, **** are p<0.05, p< 0.01, and p<0.0001 respectively. In all graphs, each dot represents one biological replicate.

To determine if *Lhx2* is enriched in the germ cells, we flow-sorted the somatic and germ cell of the fetal testis (pools of 14-16 testes) at E13.5 (3 pools) and E15.5 (2 pools). These time points were chosen because germ cells are committed to the gametogenic fate by E12.5 and have exited mitosis and committed to the spermatogenic fate by E15.5. The purity of the fractions was confirmed where the GFP+ fraction had a higher abundance of *Ddx4* (germ cell marker) with negligible amounts in the GFP-somatic cell fraction. Conversely, the somatic fraction had abundant *Amh* or *Sox9* mRNA (Sertoli cell markers) with negligible amounts in the germ cell fraction. In these fractions at E13.5 (Fig 1D) and E15.5 (Fig1E), *Lhx2* transcripts were detected in the GFP+ germ cell fraction with negligible amounts in the GFP-somatic cell fraction.

We further validated these findings using a germ cell-depleted fetal mice testis (cKIT^W^/^W^) model. RNA from WT (cKIT^+^/^+^) and germ cell-depleted testis (cKIT^W^/^W^) at E12.5 (n=3/genotype) and E15.5 (n=3/genotype) were subjected to qPCR (Fig. 1F). Results revealed that at both the time points, *Lhx2* and *Ddx4* were detected in the WT testis but negligible signals were detected in the mutant testis. These results together confirm that *Lhx2* is expressed in the germ cells of the developing testis.

*Lhx2* requires a group of co-regulators for its appropriate functions and the relative abundance of these factors in the cells is a determinant of the activity of LHX2 [27]. We observed that of the *Lhx2* co-regulators, *Ldb1* was abundantly expressed in all the cell types of developing testis, while *Ldb2* was enriched in the endothelial and interstitial lineages with lower expression in supporting cells and least in germ cells (Supplementary Figure 1B).

Of the Lim-only factors (Supplementary Figure 1B), *Lmo3* was not reported in the dataset. *Lmo1* had higher expression in the germ cells as compared to the other three cell types. *Lmo2* was enriched in endothelial cells as compared to other cell types. *Lmo4* was expressed in most somatic cells with the highest expression in supporting lineage and the lowest in the germ cells.

### Loss of *Lhx2* alters the transcriptome of the developing testis

To reveal the role of *Lhx2* in the developing testis, RNA-seq of WT and *Lhx2*^*flox/flox*^ testis at E12.5 was performed (n=30 pairs each). This time point was chosen as most cell types are committed to the male lineage in the testis. The levels of *Lhx2* mRNA was negligible in *Lhx2*^*flox/flox*^ testis while it was detected in the WT testis (Supplementary Figure 2A) indicating successful floxing of the gene. A total of 2993 genes (21% of the transcriptome) were differentially expressed (Supplementary Table 3). Amongst these, 718 (5.10%) genes were downregulated while 2275 (16.16%) genes were upregulated in *Lhx2*^*flox/flox*^ testis as compared to WT (Fig. 2A). The list of all differentially up and downregulated genes are given in Supplementary Table 4. Of the upregulated genes, 73 genes were exclusively expressed in *Lhx2*^*flox/flox*^ testis. Of the downregulated genes, 92 genes were exclusively expressed in WT testis and not detected in the *Lhx2*^*flox/flox*^ testis (Supplementary Table 4). *Ifi27l2a* gene was the maximally upregulated while *Tmem100* was the most downregulated gene in absence of *Lhx2* (Supplementary Table 4).

**Fig. 2:**
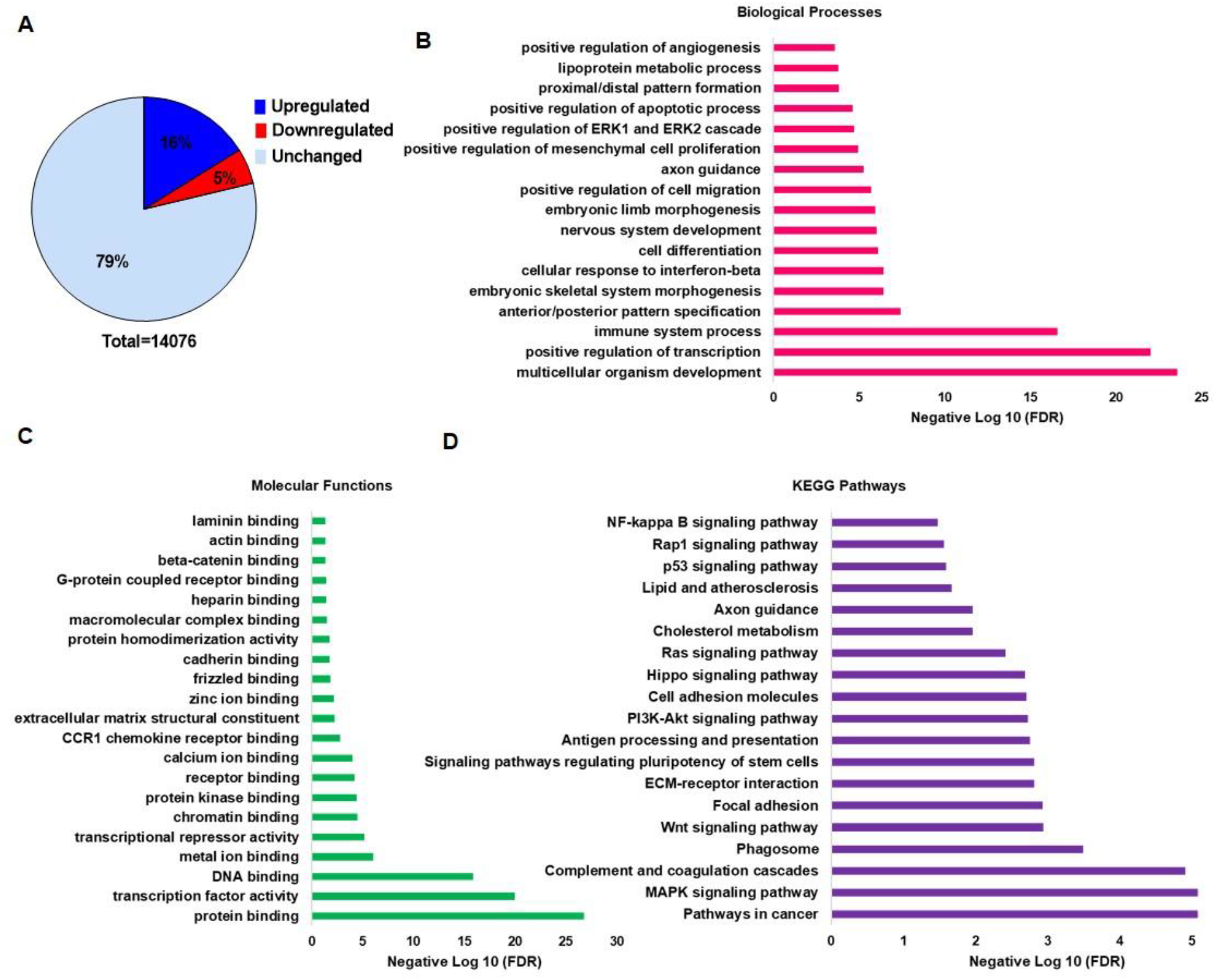
Loss of *Lhx2* alters the transcriptome of the developing testis. RNAseq for wild type (WT) and *Lhx2*^*flox/flox*^ XY embryos done at E12.5 (A) The percentage of upregulated (fold change ≥1.5) and downregulated genes (fold change ≤0.5) in the testis of *Lhx2*^*flox/flox*^ embryos as compared to WT. The false discovery rate (FDR) of the enriched biological processes (B), molecular functions (C) and, pathways (D) for the differentially expressed genes are given.

Gene Ontology analysis revealed that the differentially expressed genes were associated with biological processes like regulation of transcription, anterior/posterior patterning, cell proliferation, cell differentiation, cell migration, axon guidance, apoptosis, proximal/distal pattern formation, lipoprotein metabolic process, and angiogenesis (Fig. 2B). The associated molecular functions were protein/DNA binding, transcription factor activity, transcriptional repressor activity, chromatin binding, frizzled & β-catenin binding, ECM structural constituent, cadherin binding, GPCR binding, and actin-binding activity (Fig. 2C). In pathway analysis, there was enrichment of signaling pathways (MAPK, WNT, PI3K-Akt, Hippo, Ras, p53, Rap1, and NF-kappa B), pathways in cancer, complement and coagulation cascades, Focal/Cell adhesion, ECM-receptor interactions, and regulation of pluripotency (Fig. 2D).

### Loss of *Lhx2* in the developing XY gonads alters the transcriptome in multiple cell types

The effects of loss of *Lhx2* in the germ cells on the expression of genes enriched in different testicular cell types were examined (Supplementary Table 5). Information on 871/961 germ cell-enriched genes was available in our dataset. Nearly 13% of germ cell-enriched genes were differentially expressed, of which 8% genes were upregulated while 5% genes were downregulated (Fig. 3A). GO analysis revealed that differentially regulated genes were associated with regulation of histone methylation, regulation of transcription, and cell differentiation.

**Fig. 3:**
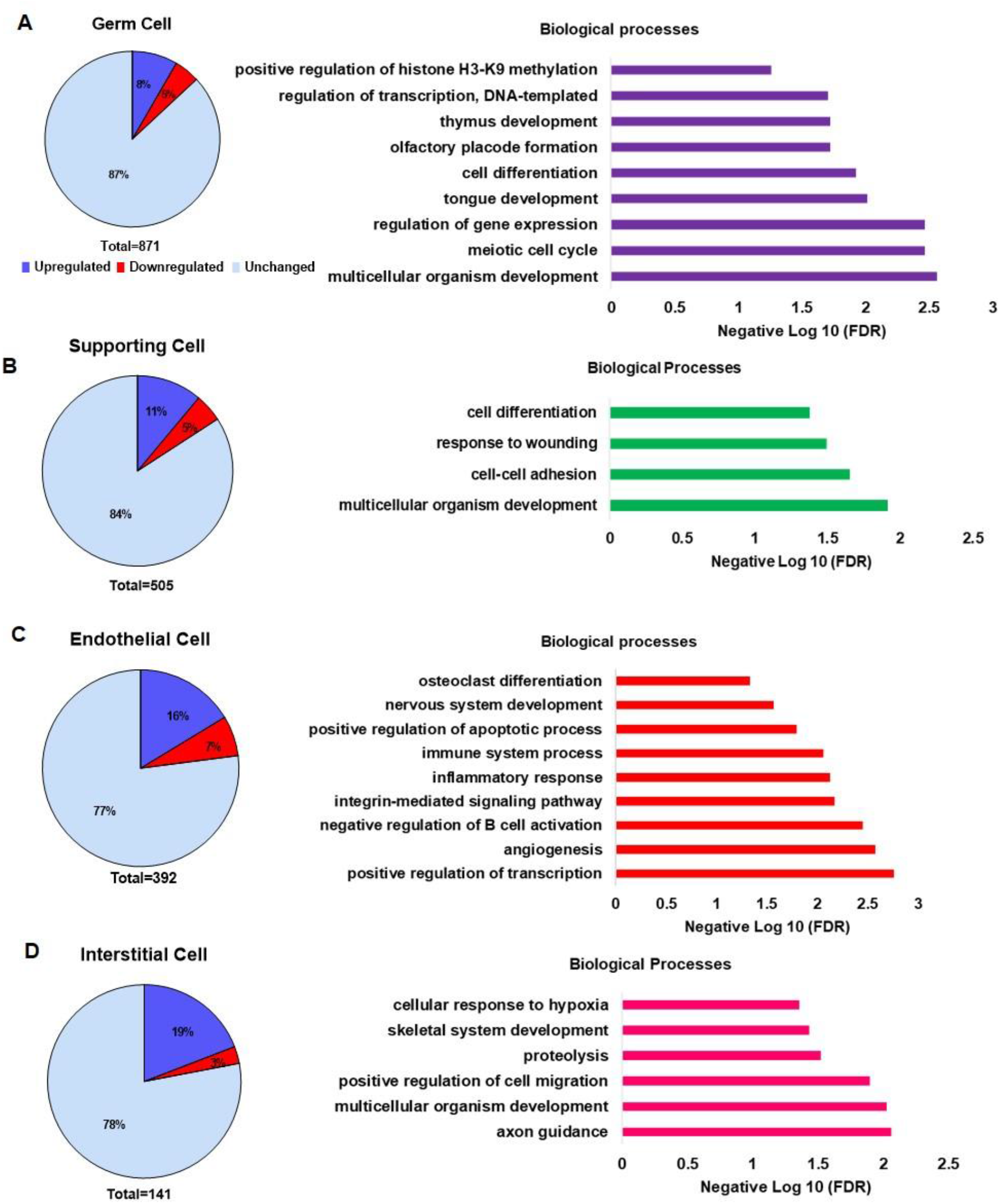
Loss of *Lhx2* in the developing XY gonads alters the transcriptome of multiple cell types. The list of genes enriched in each of the four cell types in the developing testis was extracted from the dataset GSE27715. The pie chart represents the proportion of differentially expressed genes in the Germ cells (A), Supporting cells (B), Endothelial cells (C), and Interstitial cells (D) of the testis of wild-type and *Lhx2*^*flox/flox*^ embryos at E12.5. The percentage of up and downregulated genes are given. The false discovery rate (FDR) of the enriched biological processes for the differentially expressed genes in each of the cell-types are shown as bar plots.

Of the 540 supporting lineage-enriched genes, information on 505 genes was available in our dataset. Approximately 16% of genes were differentially expressed in absence of *Lhx2*. Almost 11% of genes were upregulated and 5% were downregulated (Fig. 3B). The differentially expressed genes in the supporting lineage were associated with biological processes such as cell differentiation and cell-cell adhesion.

Of the 435 endothelial cell-enriched genes, information for 392 was available in our dataset. Nearly, 23% were differentially expressed in the *Lhx2*^*flox/flox*^ testis. Of these genes, 16% were upregulated and 7% were downregulated. The differentially expressed endothelial cell-enriched genes were associated with biological processes like angiogenesis, integrin-mediated signaling, regulation of immune response, apoptosis, nervous system development, and osteoclast differentiation (Fig. 3C).

There are 164 genes highly enriched in the interstitial cells of which information for 141 genes was available. Approximately, 22% were differentially expressed, where 19% were upregulated and 3% of genes were downregulated in *Lhx2*^*flox/flox*^ testis (Fig. 3D). These differentially expressed genes were associated with organ morphogenesis, cell migration, axon guidance, and proteolysis.

### Germ cell numbers and gametogenic fate is not compromised in *Lhx2-/-* XY gonads

The mRNA levels of the germ cell marker *Ddx4* (Fig. 4A) was similar in WT and *Lhx2-/-* testis at E13.5 (n=15 for both the genotypes), E14.5 (WT n=15, *Lhx2-/-* n=12) and E15.5 (WT n=8, *Lhx2-/-* n=6). The overall intensity of immunostained DDX4 protein was also similar in WT (n=6) and *Lhx2-/-* (n=8) XY gonads at E14.5 (Fig. 4B & C). The numbers of DDX4-positive cells and their intensity per cord were also similar in the WT and *Lhx2-/-* testis (Supplementary Figure 2B and C).

**Fig. 4:**
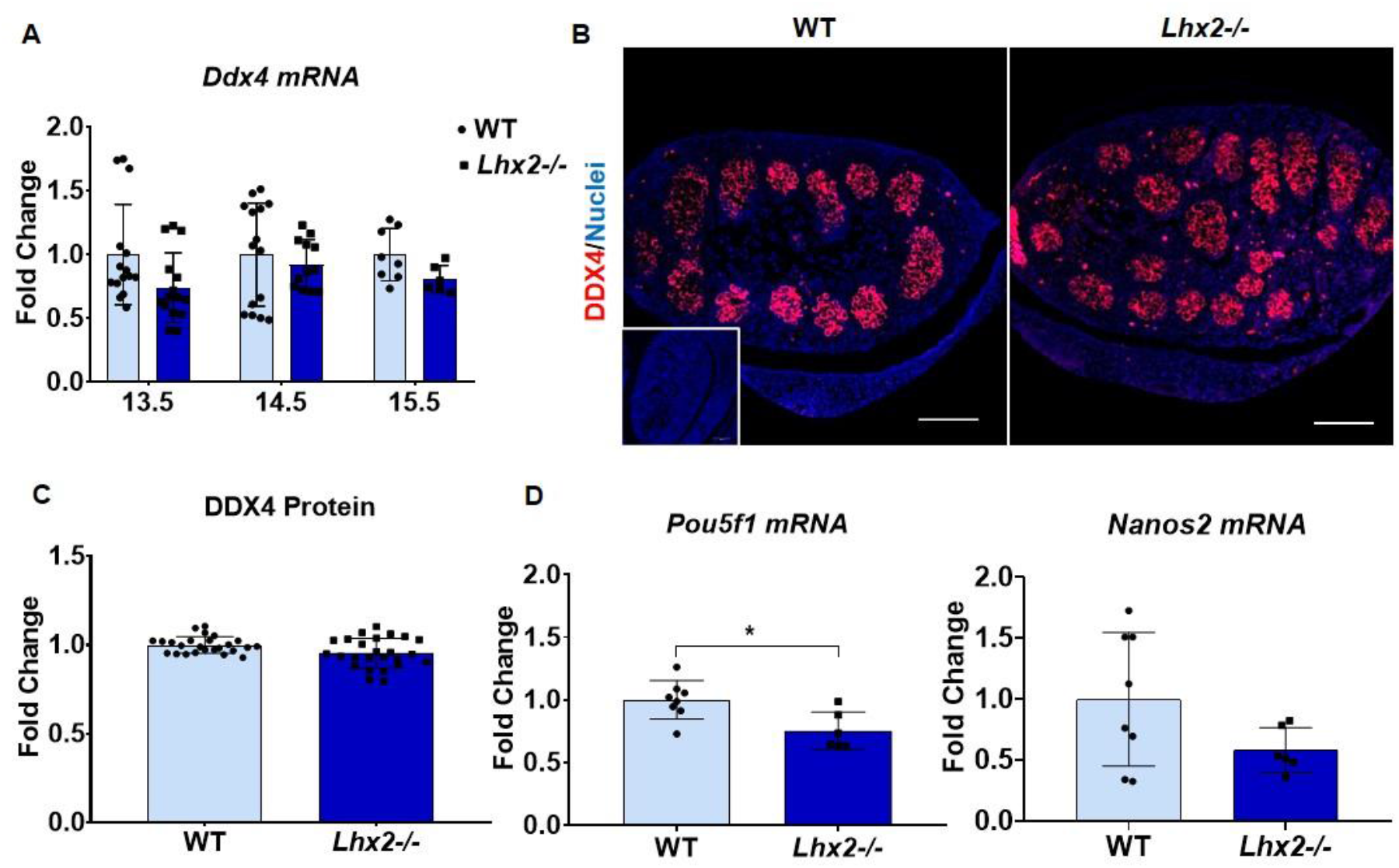
Germ cell numbers and gametogenic fate is not compromised in the developing testis in absence of *Lhx2*. (A) qPCR for *Ddx4* mRNA in wild type (WT) and *Lhx2-/-* testis at E13.5 (n=15/genotype), E14.5 (WT n=15, *Lhx2-/-* n=12) and E15.5 (WT n=8, *Lhx2-/-* n=6). (B) Immunostaining for DDX4 (germ cell marker) on sections of WT (n=6) and *Lhx2-/-* (n=8) testis at E14.5. The scale bar is 100µm. Negative control without primary antibody is shown in inset (C) Quantification of DDX4 protein intensity in WT and *Lhx2-/-* testis at E14.5. Each dot represent a single region of interest and data from different regions in multiple sections of WT (n=6) and *Lhx2-/-* (n=8) testis are plotted. (D) qPCR for *Pou5f1*, and *Nanos2* mRNA in WT (n=8) and *Lhx2-/-* (n=6) testis E15.5. Graphs are plotted as mean ± SD. Y axis is fold change where the mean value of WT was taken as 1. *Gapdh* is used for normalization in all qPCR experiments. In qPCR graphs, each dot represents one replicate. * indicates p<0.05.

The DDX4-positive germ cells are pluripotent and acquire the gametogenic fate by expressing *Dazl* and the spermatogenic fate by expressing *Nanos2* and this process is completed by E15.5 [42]. We analyzed their mRNA levels at E15.5 (WT n=8, *Lhx2-/-* n=6) and observed that *Dazl* and *Nanos2* mRNA were similar in the WT and *Lhx2-/-* testis. However, the expression of pluripotency marker *Pou5f1* was reduced significantly (p<0.05) in *Lhx2* knockout testis as compared to controls (Fig. 4D). The expression of germ cell-specific genes (*Ddx4, Pou5f1, Dazl, Piwil2*, and *Dnmt3L*) were unchanged in the WT and *Lhx2*^*flox/flox*^ testis at E12.5 (Supplementary Figure 2D). These observations imply that in absence of *Lhx2*, the number of germ cells and their commitment to gametogenic fate is not compromised though there are changes in the expression of a fraction of genes in germ cells (Fig. 3A).

### Loss of *Lhx2* does not alter Sertoli cell specification in the developing XY gonads

Loss of germ cells from developing testis due to a mutation in the cKIT gene (cKIT^W/W^) leads to a reduction in Sertoli cell number [23]. This prompted us to investigate the role of germ cell-specific *Lhx2* on the numbers and differentiation of Sertoli cells in the developing testis. Histologically, the overall testis appeared similar in *Lhx2-/-* (n=4) and WT (n=8) XY embryos (Fig. 5A). The mRNA levels of *Amh* (Sertoli cell marker) was not significantly different in the *Lhx2-/-* (n=12) and WT (n=15) testis at E14.5 (Fig. 5B) indicating that the Sertoli cells differentiated appropriately in absence of *Lhx2*. The mRNA of the female supporting cell marker, *FoxL2* was not detected in both WT (n=15) and *Lhx2-/-* (n=12) testis at E14.5 (Fig. 5C) indicating that there was no somatic cell sex reversal. Similar results were also seen at E15.5 (not shown). Somatic sex reversal was not observed at E12.5 as the expression profiles of male and female sex-determining marker genes were similar in the WT and *Lhx2*^*flox/flox*^ XY gonads (Supplementary Figure 2E). To test if Sertoli cell numbers or their differentiation was altered in absence of *Lhx2*, we profiled the expression of *Sox9*/SOX9 between E13.5 to E15.5, during the entire course of the tubular organization (Fig. 5D-G). The results revealed that levels of *Sox9* mRNA at E13.5 (WT n=15, *Lhx2-/-* n=15), E14.5 (WT n=15, *Lhx2-/-* n=12) and E15.5 (WT n=8, *Lhx2-/-* n=8) and SOX9 protein (n=6/timepoint/genotype) did not alter between WT and *Lhx2-/-* testis. However, at E15.5 the SOX9 protein intensities were marginally but significantly lower in the *Lhx2-/-* testis as compared to controls (Fig. 5D-F). Also, the number of SOX9-positive Sertoli cells per tubule was significantly reduced at both E13.5 and E15.5 (Fig. 5G). However, the total numbers of SOX9+ cells per unit area were comparable in the WT and *Lhx2-/-* testis (Supplementary Figure 2F). These results imply that *Lhx2* is dispensable for male sex determination and Sertoli cell differentiation although a proportion of the Sertoli cell transcriptome and numbers of Sertoli cells per tubule are altered (Fig. 3B).

**Fig 5:**
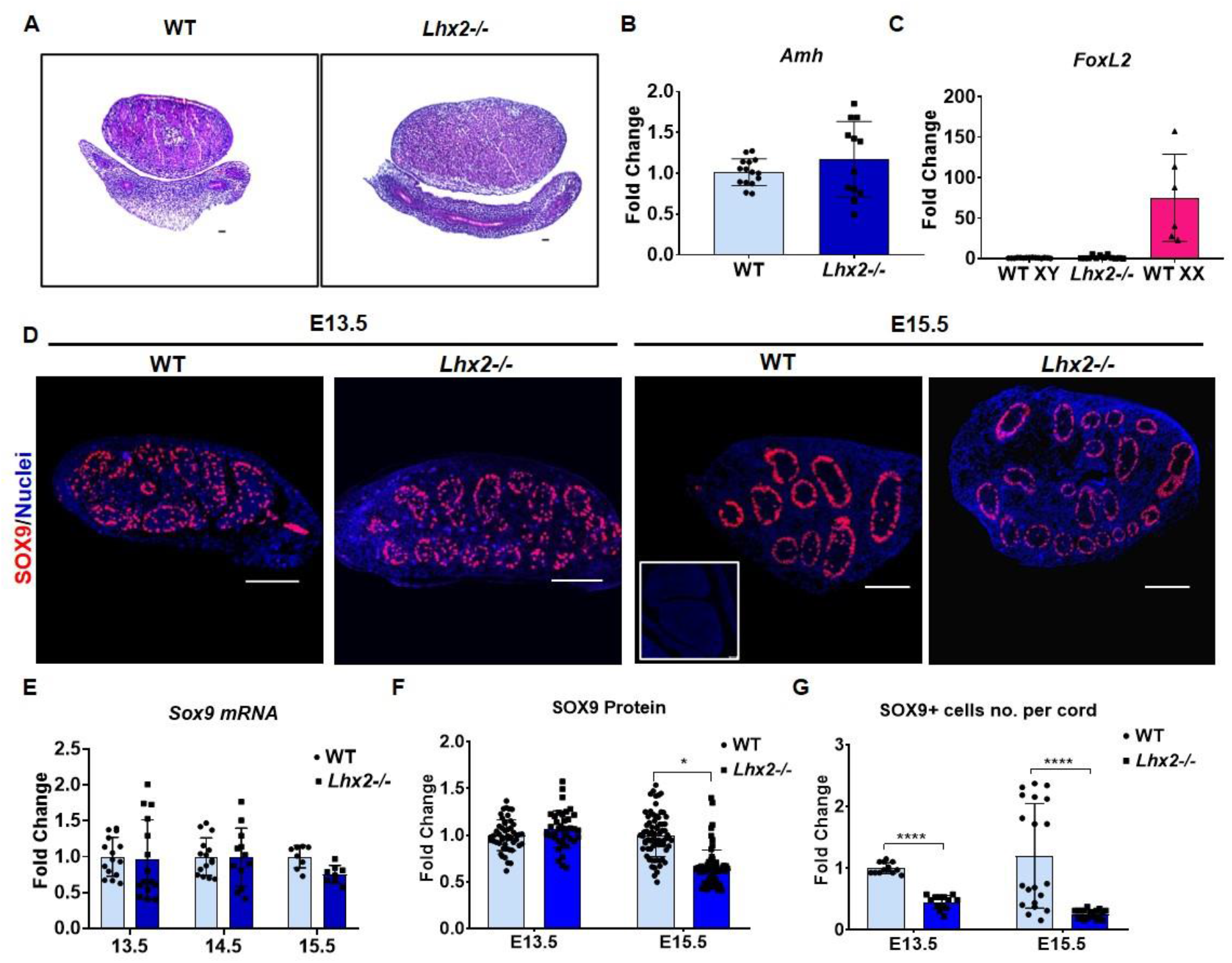
Loss of *Lhx2* does not alter Sertoli cell specification in the developing XY gonads. (A) Histology of the wild type (n=8) and *Lhx2-/-* (n=4) XY gonads at E14.5. The scale bar is 50µm. (B & C) qPCR for *Amh* (Sertoli cell marker) and *FoxL2* (granulosa cell marker) in the developing gonads at E14.5 in WT (n=15) and *Lhx2-/-* (n=12) XY gonads and WT ovaries (n=6). *Gapdh* is used for normalization in all qPCR experiments. (D) Immunostaining for SOX9 (Sertoli cell marker) in WT and *Lhx2-/-* XY at E13.5 and E15.5 (n=6/genotype/timepoint). Scale bar is 100µm. Inset is isotype control. (E) qPCR for *Sox9* mRNA at E13.5 (n=15/genotype), E14.5 (WT n=15, *Lhx2-/-* n=12) and E15.5 (n=8/genotype). Quantification of SOX9 protein (F) and SOX9 positive cell number (G) in WT and *Lhx2-/-* testis at E13.5 & E15.5 as indicated. Each dot represent a single region of interest and data from multiple independent regions in different sections of WT and *Lhx2-/-* testis are plotted. All graphs are plotted as mean ± SD. Y axis is fold change where the mean value for WT is taken as 1. In qPCR graphs, each dot represents one replicate. * and **** indicate p<0.05 and p<0.0001 respectively.

### Testicular vascularization is disrupted in absence of *Lhx2*

The observation that angiogenesis is one of the enriched biological processes in the differential transcriptome of the *Lhx2*^*flox/flox*^ testis prompted us to investigate the phenomenon further. We observed that of the 30 pro-angiogenic factors, 30% were upregulated while 10% were downregulated in absence of *Lhx2* (Fig. 6A; Supplementary Table 7) suggesting that vascularization may be altered in absence of *Lhx2*. We performed the immunoexpression of endothelial cell marker VE-Cadherin in testis sections at E13.5 when the microvasculature is organizing [43,44]. As compared to WT, there was a higher proportion of VE-Cadherin positive cells in the gonadal stroma of *Lhx2-/-* testis (n=3/genotype) (Fig. 6B). Many of these cells appeared in clusters in between the tubules indicating that endothelial cell organization is disrupted in the *Lhx2-/-* testis. The same was also reflected in whole mounts where we observed that in WT testis, microvasculature was well organized which culminated in the coelomic blood vessel; in *Lhx2-/-* testis there were hemorrhagic areas and the blood vessels were clumped in the intertubular space with no clear coelomic blood vessel (Fig. 6C). These results together indicate that testis vascularization is disrupted in absence of *Lhx2* in the germ cells.

**Fig. 6:**
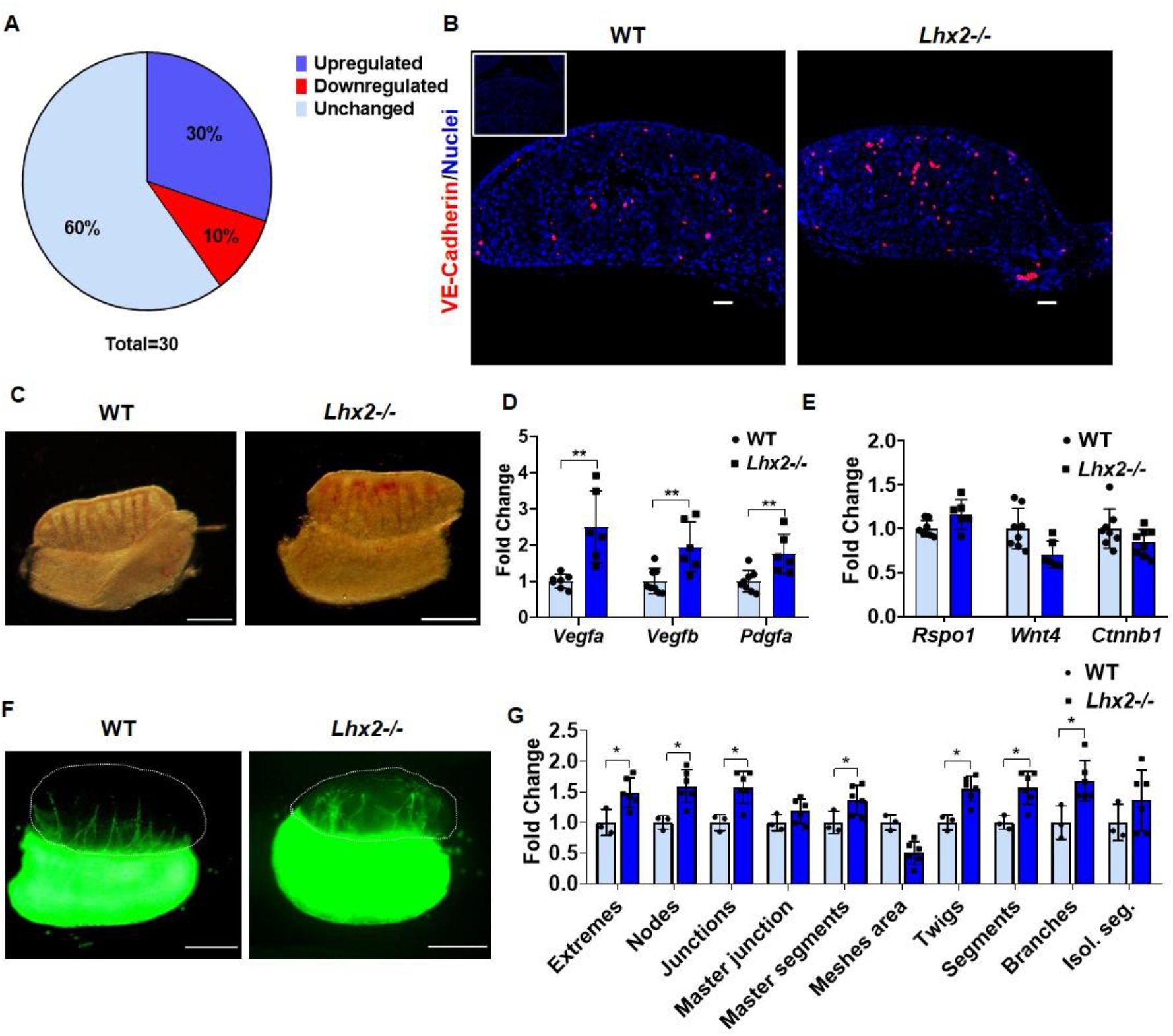
Vascularization is disrupted in the developing testis in absence of *Lhx2*. (A) The proportion of pro-angiogenic factors altered in *Lhx2*^*flox/flox*^ testis at E12.5. (B) Immunostaining of VE-cadherin (endothelial cell marker) in wild type (WT) and *Lhx2-/-* testis at E13.5 (n=3 /genotype). Scale bar is 50µm. Inset is negative control without primary antibody. (C) Brightfield image of WT and *Lhx2-/-* whole testis (n=6/genotype) at E14.5. Scale bar is 200µm qPCR for selected pro-angiogenic factors (D) and for Wnt-βcatenin genes (E) in WT (n=8) and *Lhx2-/-* (n=6) testis at E15.5. *Gapdh* is used for normalization in all qPCR experiments. (F) Gonad recombination assay of WT and *Lhx2-/-* testis overlayed on GFP (green) mesonephros from WT embryos. The recombinants were imaged 72 hrs after co-culture. Scale bar is 400µm. (G) Images were analysed using ImageJ with angiogenesis analyser plugin and the branching parameters (X axis) of the GFP-positive cells were quantitatively assessed in WT (n=3 recombinants) and *Lhx2-/-* (n=6 recombinants). All graphs are plotted as mean ± SD. Y axis is fold change where the mean value for WT is taken as 1. In all graphs, each dot represents one replicate. * and ** indicate p<0.05 and p<0.01 respectively.

Since the testicular transcriptome revealed enrichment of pro-angiogenic genes and there was disruption of testicular vascularization, we quantified the mRNA expression of pro-angiogenic factors *Vegfa, Vegfb*, and *Pdgfa* by qPCR at E15.5 when the process of testicular vascularization is completed. The results (Fig. 6D) revealed that the levels of all these genes were significantly upregulated in *Lhx2-/-* (n=6) as compared to the WT (n=8) testis. Previous studies have reported that knocking out of *Wnt4, Rspo1*, or *Ctnnb1* causes ectopic vascularization in XX gonads [45] and the overexpression of *Wnt4* disrupts vascularization in the XY gonads [46]. As Wnt/ β-catenin signaling pathways were enriched in the transcriptome *Lhx2* knockouts testis (Fig. 2), we hypothesized that altered expression of *Wnt4*/*Rspo1*/*β-catenin* may have caused the disrupted vascularization in absence of *Lhx2*. However, at E15.5, the mRNA expression of *Wnt4, Rspo1*, and *Ctnnb1* were identical in *Lhx2-/-* (n=6) and WT (n=8) testis (Fig. 6E). These results imply that disrupted angiogenesis in absence of *Lhx2* is not due to alterations in the *Wnt4, Rspo1*, and *Ctnnb1* genes but due to elevated expression of pro-angiogenic factors like *Vegfa, Vegfb*, and *Pdgfa*.

Testicular vascularization involves the migration of endothelial cells from the mesonephric vascular plexus into the gonads [8]. To investigate if the loss of *Lhx2* in the germ cells affects endothelial cell migration, we carried out gonad recombination assays and quantified the migratory pattern of endothelial cells (Fig. 6F & G). The results revealed that in the WT XY gonads (n=3), GFP-positive cells had a linear path and the cells migrated to the anterior aspect of the gonads towards the coelomic epithelium. However, in the *Lhx2-/-* gonads (n=6), GFP-positive cells had a very distorted path that deviated from their course not reaching towards the coelomic epithelium (Fig. 6F). Quantitatively, as compared to WT gonads, the measure of branching such as extremes, nodes, and junctions twigs, segments, and branches were significantly increased in absence of *Lhx2* (Fig. 6G). These results imply that the absence of *Lhx2* in the germ cells alters the intra-testicular milieu to disrupt endothelial cell migration and vascularization.

### Expansion of interstitial cells in absence of *Lhx2* in developing XY gonads

In the interstitial compartment, 22% of the enriched genes were differentially expressed and these were associated with positive regulation of cell proliferation and cell migration (Fig. 3D). To test if the interstitial compartment was altered in absence of *Lhx2*, we immunostained for the interstitial cell marker PDGFRα [47] and proliferation marker MKI67 at E14.5 when the interstitial compartment is well demarcated from the tubules. The results revealed that the PDGFRα+ cells were localized through the interstitial compartment in the wild-type testis (n=8). In the *Lhx2-/-* (n=8), PDGFRα immunoreactivity was more intense specifically in the area below the celomic epithelium (Fig. 7A). MKI67 positive cells were localized in the interstitium, their numbers appeared to be higher in the *Lhx2-/-* testis as compared to wild type (n=8/genotype). The cells below the celomic epithelium were intensely labeled with MKI67 in the *Lhx2-/-* testis (Fig. 7B). At E15.5, the mRNA levels of interstitial progenitor cell markers [47] viz *Lhx9, Nr5a1*, and *Pdgfra* were increased in *Lhx2-/-* testis (n=6) as compared to WT (n=8). The levels of the early progenitor marker *Gata4* was, however, unaltered (Fig. 7C). As compared to WT, the mRNA levels of fetal Leydig cell markers *Hoxb2*, and *Cyp17a1* were decreased in *Lhx2-/-* testis at E15.5 (Fig. 7D). The reduction in levels of *Hoxb2* but not in *Cyp17a1* was statistically significant. These results imply that in the absence of *Lhx2* there is interstitial cell expansion and defective Leydig cell differentiation

**Fig. 7:**
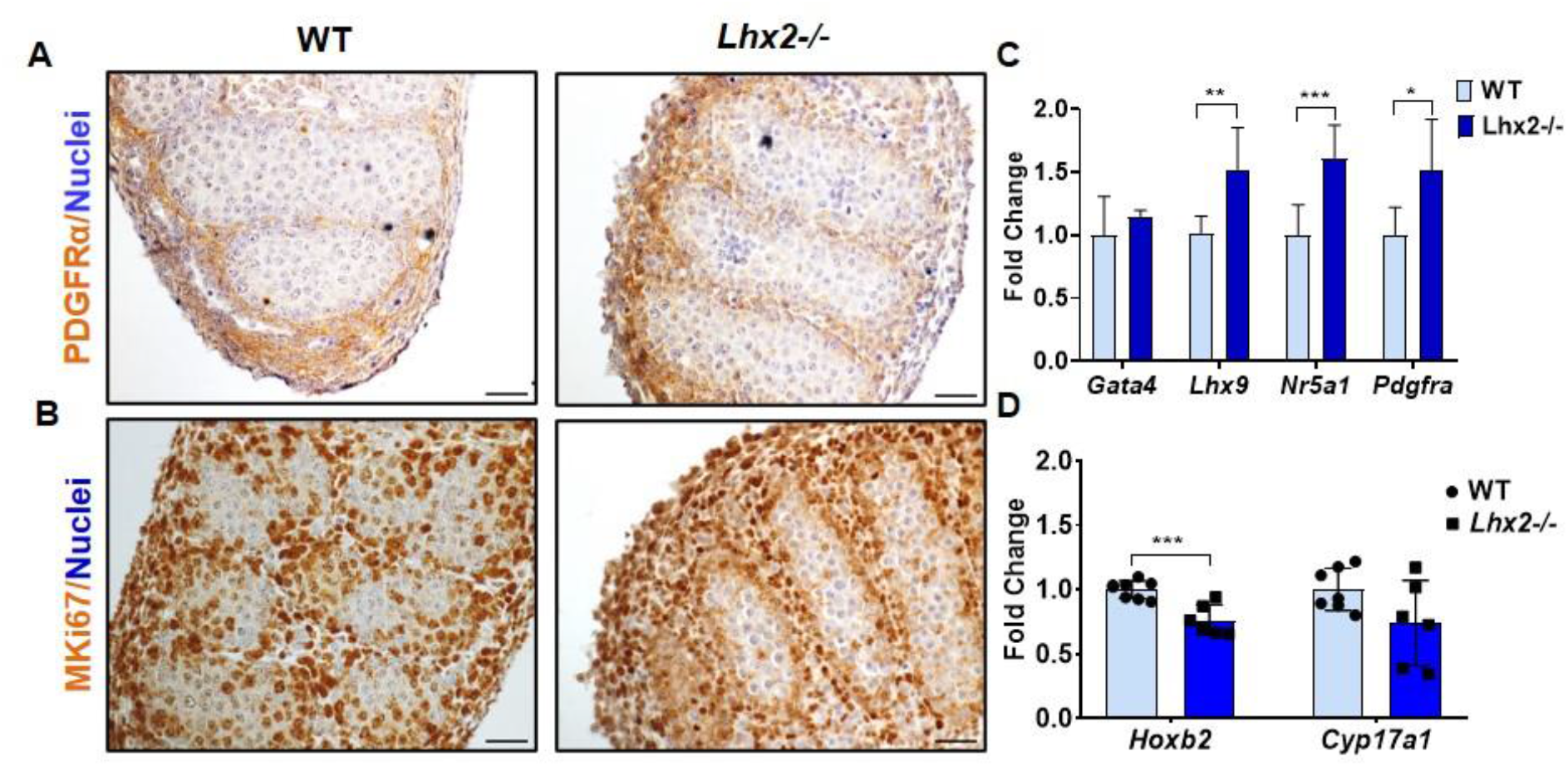
Expansion of interstitial cells in absence of *Lhx2* in the developing XY gonads. Immunostaining for (A) PDGFRα (interstitial marker) and (B) MKI67 (proliferation marker) in wild type (WT) and *Lhx2-/-* (n=8/genotype) testis at E14.5. Scale bar is 50µm. qPCR for (C) interstitial progenitors *(Gata4, Lhx9, Nr5a1*, and *Pdgfra*) and (D) fetal Leydig cell markers (*Hoxb2*, and *Cyp17a1*) in the WT (n=8) and *Lhx2-/-* (n=6) testis at E15.5. Graphs are mean ± SD. Y axis is fold change where mean value of WT was taken as 1. *Gapdh* is used for normalization in all qPCR experiments. In all graphs, each dot represents one replicate. *, ** and *** are p value<0.05, <0.01, and <0.001 respectively.

### The absence of *Lhx2* in germ cells leads to a defective tubular organization

Since there was a disruption in the supporting, endothelial and interstitial lineages (all of which aid in the tubular organization) we asked if the absence of *Lhx2* in the germ cells disrupts tubulogenesis. Towards this, co-staining of SOX9 and DDX4 was performed to visualize the testicular cords organization (Fig. 8A). As evident, the pattern of tubule organization was different in the WT (n=6) and *Lhx2-/-* testis (n=8) at E14.5. In the WT, cords were uniformly sized and had similar shapes while in *Lhx2-/-*, many tubules were abnormally shaped. Some tubules were fused, some were depleted of germ cells and others had mis-spaced Sertoli cells. The number of Sertoli cells per tubule was also reduced (Fig. 5G). Laminin staining was done to demarcate the basement membrane (Fig. 8B), and we observed that in the WT (n=5), laminin was uniformly detected at the base of the tubules. In *Lhx2-/-* testis (n=5) there was irregular laminin deposition and it did not appear well demarcated as in WT.

**Fig. 8:**
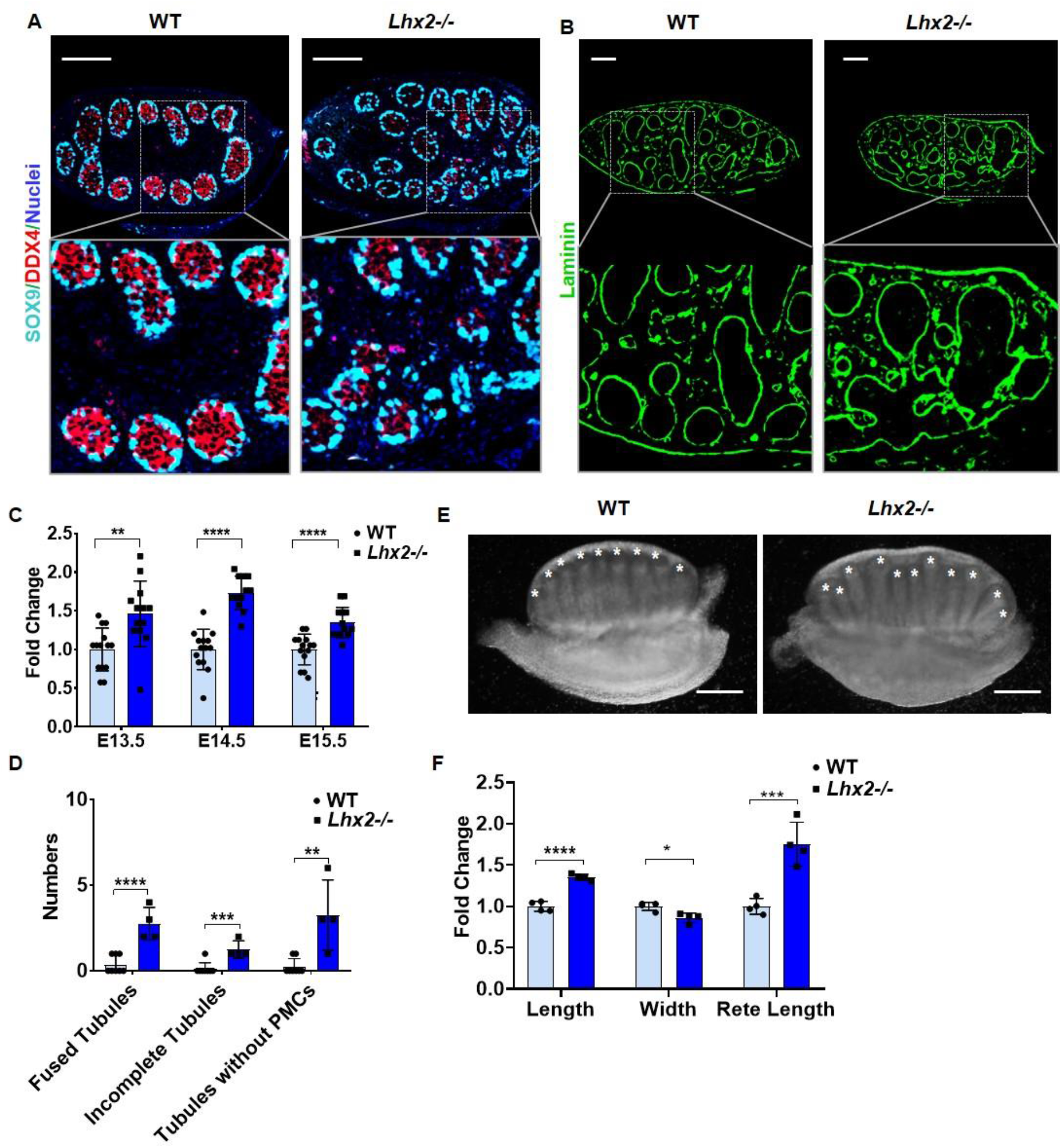
Absence of *Lhx2* leads to defect in the organization of testis cords. (A) Co-immunostaining of SOX9 and DDX4 in wild type (WT n=6) and *Lhx2-/-* (n=8) testis at E14.5. Bar is 100μm (B) Immunofluorescent detection of Laminin in the WT and *Lhx2-/-* testis (n=5/genotype) at E14.5. Scale bar is 100μm. (C) Quantification of the number of testicular cords in WT and *Lhx2-/-* testis at E13.5 (WT n=14, *Lhx2-/-* n=13), E14.5 (WT n=14, *Lhx2-/-* n=12) and E15.5 (WT n=14, *Lhx2-/-* n=13). (D) Quantification of tubular phenotypes in testicular sections of the WT (n=8) and *Lhx2-/-* (n=4) embryos at E14.5. PMCs=peritubular myoid cells. (E) Brightfield image of WT and *Lhx2-/-* whole testis at E15.5 (WT n=6, *Lhx2-/-* n=5). White asterisks mark the distal tip of cords. Scale bar is 200μm. (F) Quantification of length, width and rete length of WT and *Lhx2-/-* testis at E15.5. In all graphs, data is mean ±SD of the fold change where the mean value of WT was taken as 1. In all graphs, each dot represents one replicate and *, **, *** and **** are p value <0.05, <0.01, <0.001 and <0.0001 respectively.

We quantified the numbers of tubules per unit area in each of the histological sections of WT and *Lhx2-/-* testis at E13.5 (WT n=14, *Lhx2-/-* n=13), E14.5 (WT= n=14, *Lhx2-/-* n=12) and E15.5 (WT n=14, *Lhx2-/-* n=13) (Fig. 8C) and observed that overall there was a significant increase in the number of tubules in the *Lhx2*-/-testis as compared to WT. However, most of the tubules in the *Lhx2-/-* testis were not normal and histological analysis revealed that (Supplementary Figure 3) in some areas the testicular cords were not lined by peritubular myoid cells, many appeared fused or incomplete, and some even lacked germ cells. These phenotypes were quantified (Fig. 8D) and we observed an increase in the numbers of tubules without peritubular myoid cells layer, fused appearing tubules, and incomplete appearing tubules in the *Lhx2-/-* (n=4) testis as compared to WT (n=8). Empty tubules were rarely detected in WT, but several tubules were devoid of germ cells in *Lhx2-/-* testis. These results imply that the organization of tubules is altered in absence of *Lhx2*.

Analysis of the whole testis from WT (n=6) and *Lhx2-/-* (n=5) embryos at E15.5 (when the testis cords are completely organized), revealed that overall cords arrangements and cords length were significantly altered in *Lhx2-/-* testis (Fig. 8E). While the WT testis consistently showed well-organized tubules reaching the coelomic epithelium with the longest tubules in the center and the shortest at the poles, in the *Lhx2-/-* testis most tubules did not reach up to the coelomic region, and they were of different heights with shorter tubules interspersed between tall tubules resulting in irregular tubular arrangements. Quantification of the testis dimension (Fig. 8F) showed that as compared to WT, the length (left to right distance in the mid-region) and the rete length (left to right distance at the site of attachment to mesonephros) were significantly increased in the *Lhx2-/-*, the width (shortest diameter) was significantly reduced. These results imply that the organization of tubule and testicular dimensions are altered in absence of *Lhx2*.

## Discussion

The present study demonstrates that *Lhx2* is expressed in the germ cells of developing testis and the absence of *Lhx2* alters the endothelial and interstitial cell compartments without affecting germ cell and Sertoli cell specification resulting in the disrupted tubular organization.

*Lhx2* along with the other twelve members of the LIM-HD family and their co-factors are expressed in the developing gonads of both sexes during the window of sex determination [27]. Of these, only *Lhx1, Lhx8*, and *Lhx9* are known to have roles in the development of the reproductive system, the functions of others are not explored [27]. Previously, we have shown that *Lhx2* is expressed by the germ cells of the developing ovary [26]. Extending these findings, herein we show that *Lhx2* mRNA and protein are also expressed in the germ cells of the developing testis. Interestingly, *Ldb1* and *Lmo1* which are coregulators for *Lhx2* [48–50] are also enriched in the germ cells of the developing testis implying that the *Lhx2* co-regulator complex system may be functional in the developing germ cells.

The closest homolog of *Lhx2* is *Lhx9* which has overlapping expression patterns and functions in the brain [51]. However, in the developing gonads, *Lhx9* is expressed in somatic cells [52] while *Lhx2* is in the germ cells suggesting their divergent role in gonad development. Indeed, in the *Lhx9* knockouts, the gonad development is arrested before the onset of sex determination; the ovary [26] and the testis (present study) develop normally in absence of *Lhx2*. Also, the specification of germ cells and Sertoli cells is unaltered in *Lhx2-/-* testis, and neither the number of germ cells and Sertoli cells are compromised in *Lhx2-/-* XY gonads. Together with our previous findings [26], our results imply that *Lhx2* is dispensable for gonad development, germ cell specification, and sex determination of the supporting lineage in both sexes.

Germ cell-specific expression of *Lhx2* in developing testis intrigued us to investigate its role in testis development. RNAseq analysis revealed that the expression of almost 21% of genes are altered in the developing testis in absence of *Lhx2*. The enriched biological processes included anterior/posterior patterning, proximal/distal axis formation, limb bud morphogenesis, cell proliferation, cell differentiation, cell migration, axon guidance, and angiogenesis. The involvement of *Lhx2* in the proliferation and migration of neural stem and progenitor cells and in various cancers is well established [53,54]. *Lhx2* is involved in anterior/posterior and proximal/distal pattern formation and morphogenesis, neuron migration, and limb bud morphogenesis in many species [25,55]. Since these processes are also enriched in the testis (present study) and ovary [26] of *Lhx2* knockouts, it is tempting to propose that there would be a core transcriptome regulated by *Lhx2* which has overlapping functions irrespective of the tissue.

Since *Lhx2* is expressed by the germ cells, we first determined the changes in the expression of germ cell-enriched genes and observed that nearly 13% of them were altered. These genes were associated with germ cell development, chromatin remodeling, epithelial cell proliferation, and cell differentiation. However, the number of germ cells and their acquisition of gametogenic and spermatogenic fate is not altered in absence of *Lhx2*. These results imply that *Lhx2* is dispensable for the development and differentiation of male germ cells in the fetal testis although a proportion of its transcriptome involved in the chromatin remodeling is compromised. How this will contribute to germ cell functions needs to be investigated.

Sertoli cells are drivers of male sex determination and testicular morphogenesis [3,5]. A previous study has shown that the lack of germ cells in the developing testis affects Sertoli cell numbers and functions [23]. However, in the present study, we observed that loss of *Lhx2* does not compromise the total Sertoli cell numbers nor alters the expression of key Sertoli cell genes. To our surprise, we noticed that loss of *Lhx2* in the germ cells alters the expression of 16% of Sertoli cell-enriched genes that had functions in cell-cell adhesion and cell differentiation Indeed we did observe disruptions in the alignment of Sertoli cells of the developing testicular tubules resulting indefective tubulogenesis. These observations imply the involvement of intricate communication between the germ cells to the Sertoli cells in the developing testis.

Angiogenesis is one of the biological processes enriched in the transcriptome of *Lhx2*^*flox/flox*^ testis. These results prompted us to investigate if angiogenesis was also disrupted in the testis of *Lhx2* knockout embryos. Indeed we observed increased numbers of VE-Cadherin positive endothelial cells in the testis of *Lhx2-/-* embryos, these cells were often found in clusters and stuck in the interstitial space suggestive of disrupted vasculature. Previous studies have implicated the involvement of WNT-βcatenin pathways for angiogenesis both in the testis and ovaries [45,46,56–58]. However, we did not observe altered expression of *Wnt4, Rspo1*, and *β-catenin* in the testis of the *Lhx2-/-* embryos implying that the disrupted vasculature in the *Lhx2-/-* testis is not due to the altered WNT/β-catenin signaling. Instead, we observed that the testicular transcriptome in absence of *Lhx2* had a heightened pro-angiogenic response as there was an elevated expression of multiple angiogenesis-promoting factors and reduced expression of anti-angiogenic factors. This altered ratio of the angiogenic group of molecules disrupted the path of the migrating endothelial cells in the developing testis as evident from the gonad recombination assays. Consequently, the coelomic blood vessel was not correctly developed in the mutant testis. A similar alteration in the ratio of pro and anti-angiogenic factors and excessive endothelial cell migration was observed in the developing ovaries of *Lhx2-/-* embryos [26]. These findings imply a role of germ cell-enriched *Lhx2* in the control of endothelial cell migration in the developing testis and the ovary. In a broader sense, the results from the present study and that published earlier in the ovary [26] prompt us to propose that germ cells control the endothelial cell migration and vascularization of the developing gonads of both the sexes.

The interstitial cells of the developing testis differentiate into peritubular cells, vascular smooth muscle cells, and steroidogenic Leydig cells [58]. Absence of *Lhx2* in the germ cells altered the expression of 22% of genes enriched in the interstitial cells which are involved in cell proliferation and migration. Indeed the proliferation of coelomic epithelium was significantly increased upon loss of *Lhx2* and this expansion seems to be of the undifferentiated precursors as there was increased expression of early interstitial markers *Lhx9* and *Nr5a1* in mutant testis and the expression of the Leydig cell-enriched genes was reduced. These results imply that the germ cells control the proliferation and differentiation of the interstitial precursors in the developing testis. Whether this is due to a direct effect of the germ cell secretions or indirectly via modulation of the secretome of other cell types in the testis needs to be investigated.

In the developing testis, the organization of the tubules is governed by Sertoli cells, the correct migratory path of the endothelial cells, and the proportion of the cells in the interstitium. Since all the major cell types were affected by the loss of *Lhx2*, we investigated the tubular organization in the developing testis. To our surprise, we found that the *Lhx2-/-* testis appeared more elongated and the tubules were not appropriately organized. Also, the total numbers of tubules were higher in the *Lhx2* knockout testis; these tubules were abnormal as the numbers of SOX9 positive Sertoli cells were fewer and mis-aligned in most of the tubules. Consequently, some tubules were fused and a proportion of tubules were without the layer of peritubular myoid cells. Several tubules were also devoid of germ cells in the *Lhx2* knockout testis. In the developing testis, there is a male-specific pattern of deposition of the extracellular matrix, to ensure tubular organization. The deposition of this male-specific extracellular matrix deposition was thought to be governed by the coordinated function of Sertoli, endothelial, and interstitial cells [3,8,10,58–60]. Herein we observed that in the absence of *Lhx2* in germ cells, genes having functions in cell-cell adhesion, integrin-mediated signaling, and cell migration were differentially expressed in these three compartments and there was a disrupted organization of the basement membrane as judged by Laminin staining. These results for the first time provide conclusive evidence that germ cells govern the transcriptome of somatic cells for the organization of tubules in the developing testis. It will be of interest to study the *Lhx2-* regulated germ cell secretome to determine the mechanistic basis of this phenomenon.

To conclude, the present study has identified a crucial role of germ cells through a transcription factor, *Lhx2* in regulating the transcriptome of the supporting, vascular, and interstitial cells for governing the testicular tubule organization. The present study challenges the existing paradigm of the somatic cell-centric control of testis development and highlights an important role of germ cells in the process.

## Funding and Acknowledgement

DM lab is funded by grants from the Indian Council of Medical Research (ICMR) and the study was funded by the Department of Biotechnology (DBT) Government of India under the Grant ID: BT/PR10368/MED/97/223/2014. NS is thankful to ICMR-Senior Research Fellowship. DS is grateful to UGC-Junior Research Fellowship and AB is thankful to DBT-project Junior Research Fellowship. RS is thankful to DST-INSPIRE for the Junior and Senior Research Fellowship. We thank Prof. Shubha Tole, TIFR for providing *Lhx2*-/- and *Lhx2* floxed animals and *Lhx2* riboprobe. We thank Dr. Reddy, ICMR-NIRRCH for providing *Oct4*-GFP & cKit mice and Dr. Swanand Koli for assisting with their genotyping. We also thank NII, New Delhi for providing FVB-GFP mice. The manuscript bears the NIRRCH ID: RA/1290/07-2022

## References

[1] N. Singh, D. Modi, The Molecular Genetics of Testis Determination, in: Arafa M. et al. (Ed.), Genet. Male Infertil., Springer Cham, 2020: pp. 3–17. https://doi.org/doi.org/10.1007/978-3-030-37972-8_1.

[2] J. Cool, T. Defalco, B. Capel, Testis formation in the fetal mouse: Dynamic and complex e novo tubulogenesis, Wiley Interdiscip. Rev. Dev. Biol. 1 (2012) 847–859. https://doi.org/10.1002/wdev.62.

[3] T. Svingen, P. Koopman, Building the mammalian testis: Origins, differentiation, and assembly of the component cell populations, Genes Dev. 27 (2013) 2409–2426. https://doi.org/10.1101/gad.228080.113.

[4] H.H.-C. Yao, E. Ungewitter, H. Franco, B. Capel, Establishment of fetal Sertoli cells and their role in testis morphogenesis, Elsevier Inc., 2015. https://doi.org/10.1016/b978-0-12-417047-6.00002-8.

[5] S.P. Windley, D. Wilhelm, Signaling Pathways Involved in Mammalian Sex Determination and Gonad Development, Sex. Dev. 9 (2016) 297–315. https://doi.org/10.1159/000444065.

[6] C. Tilmann, B. Capel, Mesonephric cell migration induces testis cord formation and Sertoli cell differentiation in the mammalian gonad, Development. 126 (1999) 2883– 2890. https://doi.org/https://doi.org/10.1242/dev.126.13.2883.

[7] M. Buehr, S. Gu, A. McLaren, Mesonephric contribution to testis differentiation in the fetal mouse, Development. 117 (1993) 273–281. https://doi.org/https://doi.org/10.1242/dev.117.1.273.

[8] A.N. Combes, D. Wilhelm, T. Davidson, E. Dejana, V. Harley, A. Sinclair, P. Koopman, Endothelial cell migration directs testis cord formation, Dev. Biol. 326 (2009) 112–120. https://doi.org/10.1016/j.ydbio.2008.10.040.

[9] T. DeFalco, I. Bhattacharya, A. V. Williams, D.M. Sams, B. Capel, Yolk-sac-derived macrophages regulate fetal testis vascularization and morphogenesis, Proc. Natl. Acad. Sci. U. S. A. 111 (2014) E2384–E2393. https://doi.org/10.1073/pnas.1400057111.

[10] L. O’Donnell, L.B. Smith, D. Rebourcet, Sertoli cells as key drivers of testis function, Semin. Cell Dev. Biol. 121 (2022) 2–9. https://doi.org/10.1016/j.semcdb.2021.06.016.

[11] P. Fan, L. He, D. Pu, X. Lv, W. Zhou, Y. Sun, N. Hu, Testicular Sertoli cells influence the proliferation and immunogenicity of co-cultured endothelial cells, Biochem. Biophys. Res. Commun. 404 (2011) 829–833.https://doi.org/10.1016/j.bbrc.2010.12.068.

[12] D. Rebourcet, P.J. O’Shaughnessy, L.B. Smith, The expanded roles of Sertoli cells: lessons from Sertoli cell ablation models, Curr. Opin. Endocr. Metab. Res. 6 (2019) 42– 48. https://doi.org/10.1016/j.coemr.2019.04.003.

[13] M. Uzumcu, S.D. Westfall, K.A. Dirks, M.K. Skinner, Embryonic testis cord formation and mesonephric cell migration requires the phosphotidylinositol 3-kinase signaling pathway, Biol. Reprod. 67 (2002) 1927–1935. https://doi.org/10.1095/biolreprod.102.006254.

[14] C.A. Smith, P.J. McClive, Q. Hudson, A.H. Sinclair, Male-specific cell migration into the developing gonad is a conserved process involving PDGF signalling, Dev. Biol. 284 (2005) 337–350. https://doi.org/10.1016/j.ydbio.2005.05.030.

[15] J. Brennan, C. Tilmann, B. Capel, Pdgfr-α mediates testis cord organization and fetal Leydig cell development in the XY gonad, Genes Dev. 17 (2003) 800–810. https://doi.org/10.1101/gad.1052503.

[16] J. Cool, T.J. DeFalco, B. Capel, Vascular-mesenchymal cross-talk through Vegf and Pdgf drives organ patterning, Proc. Natl. Acad. Sci. U. S. A. 108 (2011) 167–172. https://doi.org/10.1073/pnas.1010299108.

[17] K.M. Sargent, R.M. McFee, R.S. Gomes, A.S. Cupp, Vascular endothelial growth factor A: Just one of multiple mechanisms for sex-specific vascular development within the testis?, J. Endocrinol. 227 (2015) R31–R50. https://doi.org/10.1530/JOE-15-0342.

[18] R.C. Bott, R.M. McFee, D.T. Clopton, C. Toombs, A.S. Cupp, Vascular endothelial growth factor and kinase domain region receptor are involved in both seminiferous cord formation and vascular development during testis morphogenesis in the rat, Biol. Reprod. 75 (2006) 56–67. https://doi.org/10.1095/biolreprod.105.047225.

[19] D.R. Archambeault, H.H.C. Yao, Activin A, a product of fetal Leydig cells, is a unique paracrine regulator of Sertoli cell proliferation and fetal testis cord expansion, Proc. Natl. Acad. Sci. U. S. A. 107 (2010) 10526–10531. https://doi.org/10.1073/pnas.1000318107.

[20] A.M. Clark, K.K. Garland, L.D. Russell, Desert hedgehog (Dhh) gene is required in the mouse testis for formation of adult-type Leydig cells and normal development of peritubular cells and seminiferous tubules, Biol. Reprod. 63 (2000) 1825–1838. https://doi.org/10.1095/biolreprod63.6.1825.

[21] F. Pierucci-Alves, A.M. Clark, L.D. Russell, A developmental study of the desert hedgehog-null mouse testis, Biol. Reprod. 65 (2001) 1392–1402. https://doi.org/10.1095/biolreprod65.5.1392.

[22] J.J. Meeks, S.E. Crawford, T.A. Russell, K.I. Morohashi, J. Weiss, J.L. Jameson, Dax1 regulates testis cord organization during gonadal differentiation, Development. 130 (2003) 1029–1036. https://doi.org/10.1242/dev.00316.

[23] C. Rios-Rojas, C. Spiller, J. Bowles, P. Koopman, Germ cells influence cord formation and leydig cell gene expression during mouse testis development, Dev. Dyn. 245 (2016) 433–444. https://doi.org/10.1002/dvdy.24371.

[24] S.J. Chou, S. Tole, Lhx2, an evolutionarily conserved, multifunctional regulator of forebrain development, Brain Res. 1705 (2019) 1–14. https://doi.org/10.1016/j.brainres.2018.02.046.

[25] I. Tzchori, T.F. Day, P.J. Carolan, Y. Zhao, C.A. Wassif, L.Q. Li, M. Lewandoski, M. Gorivodsky, P.E. Love, F.D. Porter, H. Westphal, Y. Yang, LIM homeobox transcription factors integrate signaling events that control three-dimensional limb patterning and growth, Development. 136 (2009) 1375–1385. https://doi.org/10.1242/dev.026476.

[26] N. Singh, D. Singh, A. Bhide, R. Sharma, S. Sahoo, M.K. Jolly, D. Modi, Lhx2 in germ cells suppresses endothelial cell migration in the developing ovary, Exp. Cell Res. 415 (2022) 113108. https://doi.org/10.1016/J.YEXCR.2022.113108.

[27] N. Singh, D. Singh, D. Modi, LIM Homeodomain (LIM-HD) Genes and Their Co-Regulators in Developing Reproductive System and Disorders of Sex Development, Sex. Dev. (2021) 1–15. https://doi.org/10.1159/000518323.

[28] N. Singh, D. Singh, A. Bhide, R. Sharma, S. Bhowmick, V. Patel, D. Modi, LHX2 in germ cells control tubular organization in the developing mouse testis, BioRxiv. (2022) 1–38. https://doi.org/https://doi.org/10.1101/2022.12.29.522214.

[29] T. Yoshimizu, N. Sugiyama, M. De Felice, Y. Yeom, K. Ohbo, K. Masuko, M. Obinata, A. Kuniya, H.R. Schöler, Y. Matsui, Germline-specific expression of the Oct-4/green fluorescent protein (GFP) transgene in mice, Dev. Growth Differ. 41 (1999) 675–684. https://doi.org/10.1046/j.1440-169X.1999.00474.x.

[30] C. Waskow, S. Paul, C. Haller, M. Gassmann, H.R. Rodewald, Viable c-KitW/W mutants reveal pivotal role for c-Kit in the maintenance of lymphopoiesis, Immunity. 17 (2002) 277–288. https://doi.org/10.1016/S1074-7613(02)00386-2.

[31] F.D. Porter, J. Drago, Y. Xu, S.S. Cheema, C. Wassif, S.P. Huang, E. Lee, A. Grinberg, J.S. Massalas, D. Bodine, F. Alt, H. Westphal, Lhx2, a LIM homeobox gene, is required for eye, forebrain, and definitive erythrocyte development, Development. 124 (1997) 2935–2944. https://doi.org/https://doi.org/10.1242/dev.124.15.2935.

[32] V.S. Mangale, K.E. Hirokawa, P.R.V. Satyaki, N. Gokulchandran, S. Chikbire, L. Subramanian, A.S. Shetty, B. Martynoga, J. Paul, M. V. Mai, Y. Li, L.A. Flanagan, S. Tole, E.S. Monuki, Lhx2 selector activity specifies cortical identity and suppresses hippocampal organizer fate, Science (80-.). 319 (2008) 304–309. https://doi.org/10.1126/science.1151695.

[33] G. Godbole, D. Modi, Regulation of decidualization, interleukin-11 and interleukin-15 by homeobox A 10 in endometrial stromal cells, J. Reprod. Immunol. 85 (2010) 130– 139. https://doi.org/10.1016/j.jri.2010.03.003.

[34] D. Modi, S. Sane, D. Bhartiya, Over-expression of müllerian inhibiting substance mRNA in the turner syndrome ovary, Sex. Dev. 3 (2009) 245–252. https://doi.org/10.1159/000261659.

[35] G. Godbole, A. Roy, A.S. Shetty, S. Tole, Novel functions of LHX2 and PAX6 in the developing telencephalon revealed upon combined loss of both genes, Neural Dev. 12 (2017) 1–8. https://doi.org/10.1186/s13064-017-0097-y.

[36] S. Bulchand, E.A. Grove, F.D. Porter, S. Tole, LIM-homeodomain gene Lhx2 regulates the formation of the cortical hem, Mech. Dev. 100 (2001) 165–175. https://doi.org/10.1016/S0925-4773(00)00515-3.

[37] L. Wojtasz, K. Daniel, A. Toth, Fluorescence activated cell sorting of live female germ cells and somatic cells of the mouse fetal gonad based on forward and side scattering, Cytom. Part A. 75 (2009) 547–553. https://doi.org/10.1002/cyto.a.20729.

[38] A. Mishra, M. Galvankar, N. Singh, D. Modi, Spatial and temporal changes in the expression of steroid hormone receptors in mouse model of endometriosis, J. Assist. Reprod. Genet. 37 (2020) 1069–1081. https://doi.org/10.1007/s10815-020-01725-6.

[39] A. Mishra, M. Galvankar, S. Vaidya, U. Chaudhari, D. Modi, Mouse model for endometriosis is characterized by proliferation and inflammation but not epithelial-to-mesenchymal transition and fibrosis, J. Biosci. 45 (2020) 105. https://doi.org/10.1007/s12038-020-00073-y.

[40] A. Tiwari, N. Ashary, N. Singh, S. Sharma, D. Modi, Modulation of E-Cadherin and N-Cadherin by ovarian steroids and embryonic stimuli, Tissue Cell. 73 (2021) 101670. https://doi.org/10.1016/j.tice.2021.101670.

[41] S.A. Jameson, A. Natarajan, J. Cool, T. DeFalco, D.M. Maatouk, L. Mork, S.C. Munger, B. Capel, Temporal transcriptional profiling of somatic and germ cells reveals biased lineage priming of sexual fate in the fetal mouse gonad, PLoS Genet. 8 (2012) e1002575. https://doi.org/10.1371/journal.pgen.1002575.

[42] R. Saba, Y. Kato, Y. Saga, NANOS2 promotes male germ cell development independent of meiosis suppression, Dev. Biol. 385 (2014) 32–40. https://doi.org/10.1016/j.ydbio.2013.10.018.

[43] D. Coveney, J. Cool, T. Oliver, B. Capel, Four-dimensional analysis of vascularization during primary development of an organ, the gonad, Proc. Natl. Acad. Sci. U. S. A. 105 (2008) 7212–7217. https://doi.org/10.1073/pnas.0707674105.

[44] and P.K. N, Combes AAlexander N. Combes1, Emmanuelle Lesieur1, Vincent R. Harley2, Andrew H. Sinclair3, Melissa H. Little1, Dagmar Wilhelm1, Three-dimensional visualisation of testis cord morphogenesis, a novel tubulogenic mechanism in development, Dev Dyn. 238 (2009) 1033–1041. https://doi.org/10.1002/dvdy.21925.Three-dimensional.

[45] A.A. Chassot, I. Gillot, M.C. Chaboissier, R-spondin1, WNT4, and the ctnnb1 signaling pathway: Strict control over ovarian differentiation, Reproduction. 148 (2014) R97– R110. https://doi.org/10.1530/REP-14-0177.

[46] B.K. Jordan, J.H.C. Shen, R. Olaso, H.A. Ingraham, E. Vilain, Wnt4 overexpression disrupts normal testicular vasculature and inhibits testosterone synthesis by repressing steroidogenic factor 1/β-catenin synergy, Proc. Natl. Acad. Sci. U. S. A. 100 (2003) 10866–10871. https://doi.org/10.1073/pnas.1834480100.

[47] L. Garcia-Alonso, V. Lorenzi, C.I. Mazzeo, J.P. Alves-Lopes, K. Roberts, C. Sancho-Serra, J. Engelbert, M. Marečková, W.H. Gruhn, Rachel A. Botting, T. Li, B. Crespo, Stijn van Dongen, V.Y. Kiselev, E. Prigmore, M. Herbert, A. Moffett, A. Chédotal, O.A. Bayraktar, A. Surani, M. Haniffa, R. Vento-Tormo, Gonadal, Single-cell roadmap of human gonadal development, Nature. 607 (2022) 540–547. https://doi.org/10.1038/s41586-022-04918-4.

[48] V. Kinare, A. Iyer, H. Padmanabhan, G. Godbole, T. Khan, Z. Khatri, U. Maheshwari, B. Muralidharan, S. Tole, An evolutionarily conserved Lhx2-Ldb1 interaction regulates the acquisition of hippocampal cell fate and regional identity, Development. 147 (2020) 1–6. https://doi.org/10.1242/dev.187856.

[49] V. Lakhina, L. Subramanian, D. Huilgol, A.S. Shetty, V.A. Vaidya, S. Tole, Seizure evoked regulation of LIM-HD genes and co-factors in the postnatal and adult hippocampus, F1000Research. 2 (2013) 1–12. https://doi.org/10.12688/f1000research.2-205.v1.

[50] J. De Melo, B.S. Clark, A. Venkataraman, F. Shiau, C. Zibetti, S. Blackshaw, Ldb1-and Rnf12-dependent regulation of Lhx2 controls the relative balance between neurogenesis and gliogenesis in the retina, Development. 145 (2018) dev159970. https://doi.org/10.1242/dev.159970.

[51] D. Peukert, S. Weber, A. Lumsden, S. Scholpp, Lhx2 and Lhx9 determine neuronal differentiation and compartition in the caudal forebrain by regulating Wnt signaling, PLoS Biol. 9 (2011) e1001218. https://doi.org/10.1371/journal.pbio.1001218.

[52] O.S. Birk, D.E. Caslano, C.A. Wassif, T. Cogilatl, L. Zhaos, Y. Zhao, A. Grinberg, S.P. Huang, J.A. Kreidberg, K.L. Parker, F.D. Porter, H. Westphal, The LIM homeobox gene Lhx9 is essential for mouse gonad formation, Nature. 403 (2000) 909–913. https://doi.org/10.1038/35002622.

[53] R.F. Hevner, Evolution of the mammalian dentate gyrus, J. Comp. Neurol. 524 (2016) 578–594. https://doi.org/10.1002/cne.23851.

[54] X. Li, X. Wu, H. Chen, Z. Liu, H. He, L. Wang, LHX2 Enhances the Malignant Phenotype of Esophageal Squamous Cell Carcinoma by Upregulating the Expression of SERPINE2, Genes (Basel). 13 (2022) 1457–1469. https://doi.org/10.3390/genes13081457.

[55] S.J. Chou, D.D.M. O’Leary, Role for Lhx2 in corticogenesis through regulation of progenitor differentiation, Mol. Cell. Neurosci. 56 (2013) 1–9. https://doi.org/10.1016/j.mcn.2013.02.006.

[56] S.G. Tevosian, N.L. Manuylov, To β or not to β: Canonical β-catenin signaling pathway and ovarian development, Dev. Dyn. 237 (2008) 3672–3680. https://doi.org/10.1002/DVDY.21784.

[57] K. Jeays-Ward, M. Dandonneau, A. Swain, Wnt4 is required for proper male as well as female sexual development, Dev. Biol. 276 (2004) 431–440. https://doi.org/10.1016/j.ydbio.2004.08.049.

[58] A. Heinrich, T. Defalco, Essential roles of interstitial cells in testicular development and function, Andrology. 8 (2020) 903–914. https://doi.org/10.1111/andr.12703.

[59] P.S. Tung, M.K. Skinner, I.B. Fritz, Cooperativity between Sertoli Cells and Peritubular Myoid Cells in the Formation of the Basal Lamina in the Seminiferous Tubule, Ann. N. Y. Acad. Sci. 438 (1984) 435–446. https://doi.org/10.1111/j.1749-6632.1984.tb38304.x.

[60] S.R. Chen, Y.X. Liu, Testis cord maintenance in mouse embryos: Genes and signaling, Biol. Reprod. 94 (2016) 1–7. https://doi.org/10.1095/biolreprod.115.137117.

