## SUPPLEMENTARY FIGURE 1-3 for "LHX2 in germ cells control tubular organization in the developing mouse testis"

### Supplementary Figures

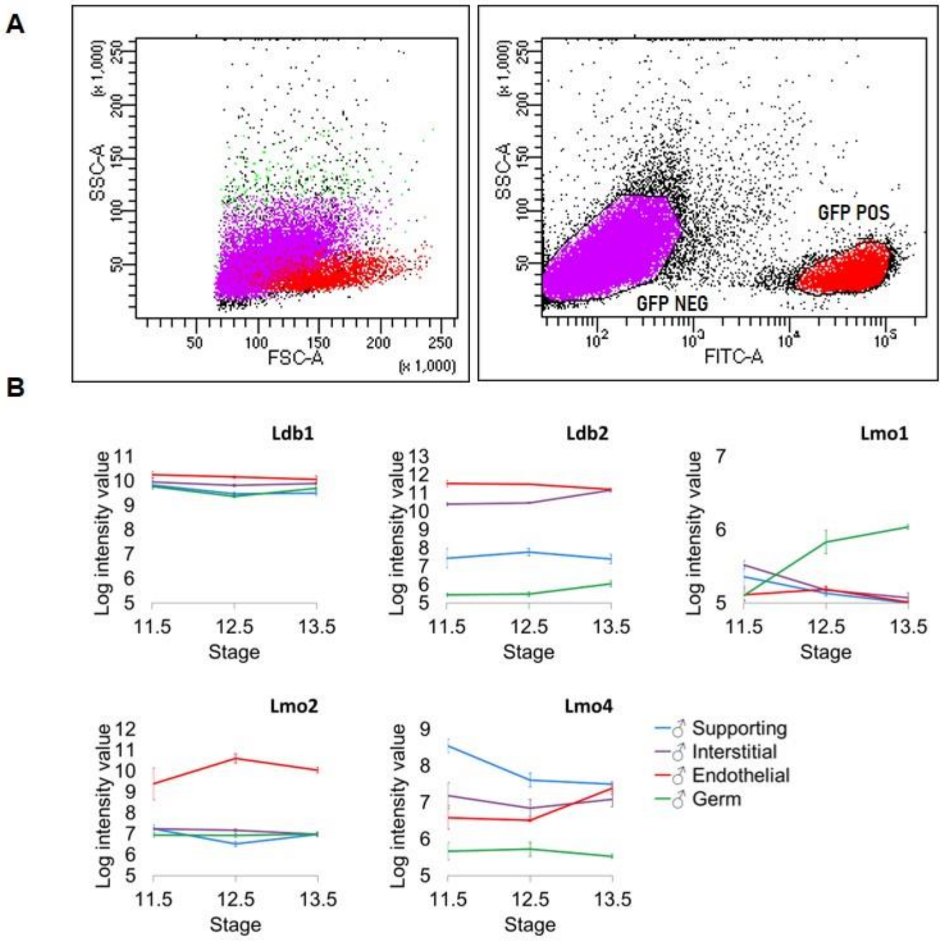


**Supplementary Fig 1:** (A) Gating (forward and side scatter) criteria and GFP-based sorting of germ and somatic cells in the developing testis. (B) Expression atlas of LIM-HD co-factors (LMOs and LDBs) in different cell types of developing testis at E11.5-13.5. Graph is plotted from data in GSE27715. The Y axis is log-transformed, normalized intensity values and the X axis shows the developmental time points.

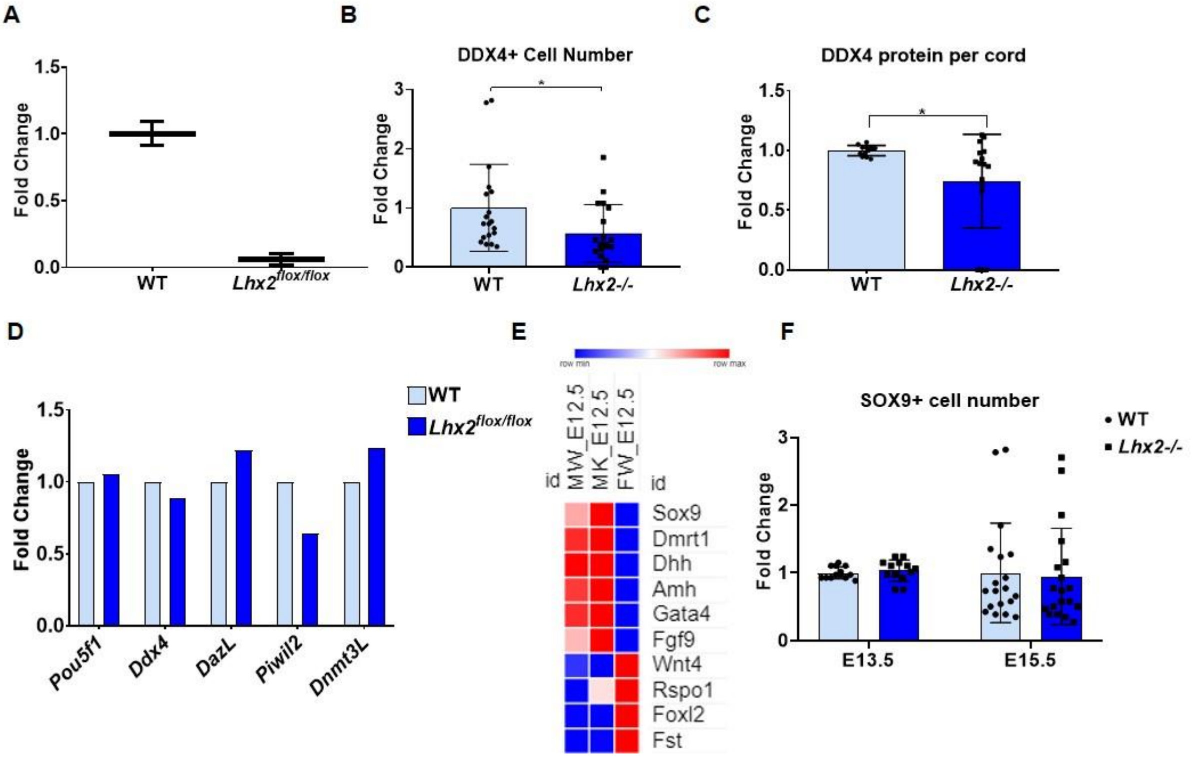


**Supplementary Fig 2: (**A) Expression of *Lhx2* in wild type (WT) and *Lhx2$^{flox/flox}$* testis at E12.5. Quantification of DDX4 positive cells per cord (B) and DDX4 protein per cord (C) in WT (n=6) and *Lhx2-/-* (n=8) testis at E14.5. Y axis is fold change where the mean of WT is taken as 1. * is $p< 0.05$. (D) Expression of germ cell-specific genes in WT and *Lhx2$^{flox/flox}$* testis at E12.5. FPKM values were extracted from RNAseq data. Fold change was calculated where the value for WT was taken as 1. (E) Heatmap of male and female sex-determining genes in the wild type XY (MW), XX (FW), and *Lhx2-/-* XY (MK) gonads at E12.5. Data is extracted from the RNAseq analysis and normalized by row minimum (F) Quantification of SOX9 positive cells per unit area in WT and *Lhx2-/-* testis at E13.5 and E15.5. Y axis is fold change where the mean of WT is taken as 1.

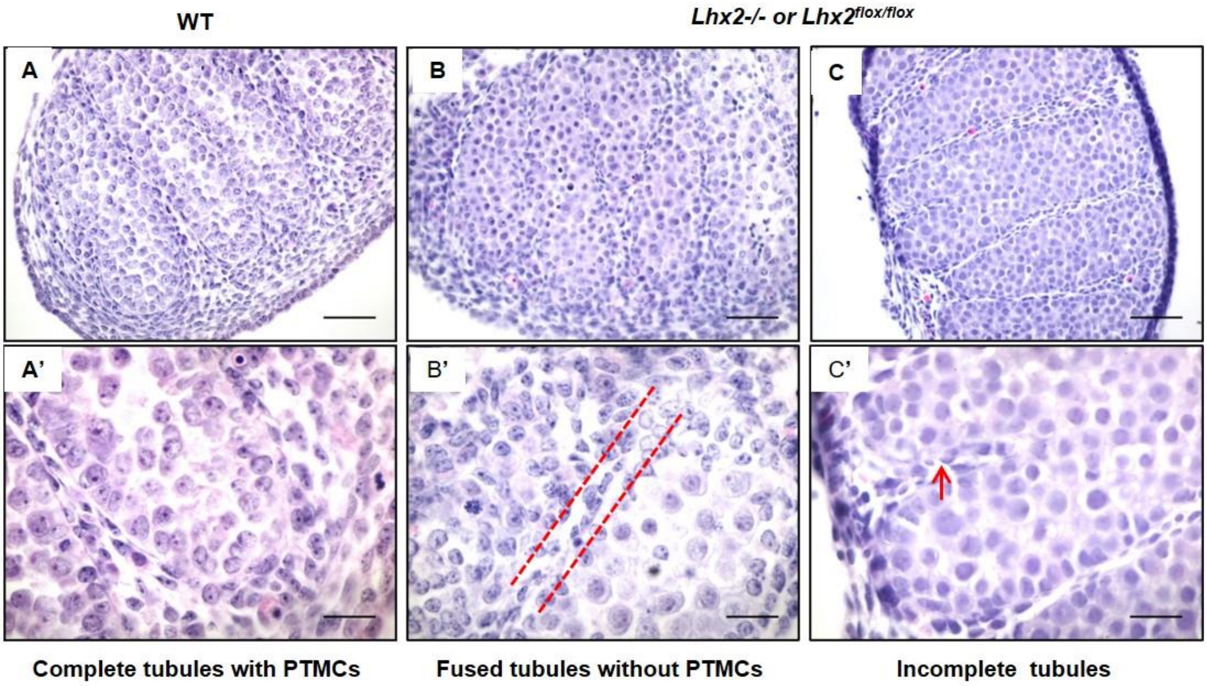


**Supplementary Fig 3:** Histology of wild type (WT) (panel A) and *Lhx2* knockout testis (*Lhx2*-/- or *Lhx2*$^{flox/flox}$, panel B and C) at E14.5 showing various defects of tubule organization. The upper panel (A-C) is low magnification images (Scale bar = 50µm) and the lower panel (A'-C') is the higher magnification (scale bar = 20µm). The red parallel lines show the absence of peritubular myoid cells between the two cords. The red arrow shows an incomplete tubule.
