## SUPPLEMENTARY TABLE 1 for "LHX2 in germ cells control tubular organization in the developing mouse testis"

**Supplementary Table.1: Sequences, optimized annealing temperature and amplicon size of the primers used in the study.**

Primers with * marks are used for genotyping

| **Gene** | **Sequence (5’ to 3’)** | **Optimized annealing temperature** | **Amplicon**  **Size** |
| --- | --- | --- | --- |
| *Gapdh* | FP-5’ GGCCGGGGCCCACTTGAAG 3’  RP-5’ TGGATGACCTTGGCCAGGGGG 3’ | 68^o^C | 174 bps |
| *Lhx2* | FP-5’ CCTACCCCAGCAGCCAAAAG 3’  RP-5’ CTGGAGGACTCTCTTGGTGAG 3’ | 61^o^C | 158 bps |
| *Amh* | FP-5’ GCGGGGCACACAGAACCTCTG 3’  RP-5’ TCCGCGTGAAACAGCGGGAA 3’ | 64^o^C | 191 bps |
| *Ddx4* | FP-5’ CCTGGAGCGGAGAGGAACCTGAA 3’  RP-5’ TTCCAGAAGGCCCATCTTCCATTTC 3’ | 67^o^C | 183 bps |
| *Oct4/Pou5f1* | FP-5’ AGCTGCTGAAGCAGAAGAGG 3’  RP-5’ GGTTCTCATTGTTGTCGGCT 3’ | 64^o^C | 198 bps |
| *Nanos2* | FP-5’AGTCTCTCTACCGACGCAGT 3’  RP-5’AAACGTTGAGATCGGGGGAC 3’ | 62^o^C | 168 bps |
| *Sox9* | FP-5’ CACAAGAAAGACCACCCCGA 3’  RP-5’ GGACCCTGAGATTGCCCAGA 3’ | 68^o^C | 209 bps |
| *Foxl2* | FP-5’ GGCTCTTCGGGAGCGGAGGA 3’  RP-5’ TGGCAGGAGGCGTAGGGCAT 3’ | 61^o^C | 163 bps |
| *Vegfa* | FP-5’ GGACGGGCCTCCGAAACCATG 3’  RP-5’ GACGGCAGTAGCTTCGCTGGT 3’ | 62^o^C | 175 bps |
| *Vegfb* | FP-5’ TGACATCATCCATCCCACTC 3’  RP-5’ CCTTGGCAATGGAGGAAG 3’ | 58^o^C | 125 bps |
| *Pdgfa* | FP-5’ GAGGAAGCCGAGATACCCC 3’  RP-5’ GGCACATGGTTAATGGCATGG 3’ | 58^o^C | 164 bps |
| *Rspo1* | FP-5’ GCCGCTGCGCCAGGTCTATC 3’  RP-5’ AGAGCCAGGCCCGGATCCAC 3’ | 63^o^C | 175 bps |
| *Wnt4* | FP-5’ TGGACTCCCTCCCTGTCTTTGGGA 3’  RP-5’ TCCTGACCACTGGAAGCCCTGTG 3’ | 64^o^C | 188 bps |
| *Ctnnb1* | FP-5’ GCGGCCGCGAGGTACCTGAA 3’  RP-5’ GAAGGAGCTGTGGTGGTGGCA 3’ | 60^o^C | 192 bps |
| *Gata4* | FP-5’ CCCTGGAAGACACCCCAATC 3’  RP-5’ AAGCGGCAGGCGGCGCT 3’ | 57^o^C | 199 bps |
| *Lhx9* | FP-5’ GGACCTCAAACAGCTTGCTC 3’  RP-5’ AATTTTCAAACGTCGGGATG 3’ | 60^o^C | 103 bps |
| *Nr5a1* | FP-5’ GCTTGAATGGCACTCTACGG 3’  RP-5’ AGTTGTAGACATGAGAGACGGTG 3’ | 62^o^C | 341 bps |
| *Pdgfra* | FP-5’ GAGCGTGCTAGGGCGGAACC 3’  RP-5’ CCCGGCCCTGTGAGGAGACA 3’ | 66^o^C | 167 bps |
| *Hoxb2* | FP-5’CACCATTGAAAGCCATGAATT  RP-5’GGAGGAATTAATTGTCGACTCCTT | 54 ^o^C | 149 bps |
| *Cyp17a1* | FP-5’ GATCGGTTTATGCCTGAGCG 3’  RP-5’ CATGGGATCCGGGACGTTAG 3’ | 67 ^o^C | 329 bps |
| *Jarid** | FP-5’ CTGAAGCTTTTGGCTTTGAG 3’  RP-5’ CCACTGCCAAATTCTTTGG 3’ | 61^o^C | 331 bps  302 bps |
| *Lhx2* WT* | FP-5’ ACCAGACTCAGGGGAAACTCAG 3’  RP-5’ GTGACTGAACTCCGAACCATTG 3’ | 61^o^C | 385 bps |
| *Lhx2* Null* | FP-5’ ACCAGACTCAGGGGAAACTCAG 3’  RP-5’ ATGCCTGCTTGCCGAATATC 3’ | 61^o^C | 550 bps |
| *cKit******** | FP-5’ TTCCTTGCAGAGCAAATCCAG 3’  RP-5’ ATACATGGGTTTCTGGAGGAG 3’ | 61^o^C | 231 bps |
