## SUPPLEMENTARY TABLE 2 for "LHX2 in germ cells control tubular organization in the developing mouse testis"

**Supplementary Table. 2 list of antibodies, their sources and optimized concentrations used in this study**

| **Protein** | **Catalogue No** | **Host** | **Working dilution** |
| --- | --- | --- | --- |
| LHX2 | Santacruz (C-20) (sc-19344) | Goat | 1:400 |
| SOX9 | Santacruz (H-90) (sc-20095) or Novus Biologicals, (NBP1-85551) | Rabbit | 1:100 |
| VASA | Novus Biologicals, (NBP2-24558) | Rabbit | 1:100 |
| PDGFRα | Abcam (ab61219) | Rabbit | 1:100 |
| MKI67 | Novus Biologicals, (NB110-89717) | Rabbit | 1:1500 |
| Laminin | Novus Biologicals, (NB300-144) | Rabbit | 1:200 |
