## SUPPLEMENTARY TABLE 3 for "LHX2 in germ cells control tubular organization in the developing mouse testis"

**Supplementary Table. 3 Numbers of differentially expressed genes in *Lhx2^flox/flox^* testis at E12.5 as revealed by RNAseq analysis**

| **Lhx2^flox/flox^ vs WT** | |
| --- | --- |
| **Regulation** | **Number of Genes (%)** |
| Upregulated (≥ 1.5 fold) | 2275 (16.16%) |
| Downregulated (≤ 0.5 fold) | 718 (5.1%) |
| Unchanged | 11083 (78.73%) |
