## SUPPLEMENTARY TABLE 5 for "LHX2 in germ cells control tubular organization in the developing mouse testis"

**Supplementary Table. 5 Differentially expressed genes in different cell types of *Lhx2^flox/flox^* testis at E12.5 as identified by deconvolution of the RNAseq data**

| **Cell Type** | **Upregulated genes**  **Numbers (% of total genes)** | **Downregulated**  **Numbers (% of total genes)** |
| --- | --- | --- |
| Germ cells | 73 (8.38%) | 41 (4.70%) |
| Supporting cells | 56 (11.08%) | 24 (4.75%) |
| Endothelial cells | 64 (16.32%) | 26 (6.63%) |
| Interstitial cells | 27 (19.14%) | 4 (2.83%) |
